# RNA-sequencing manifests the intrinsic role of MAPKAPK2 in facilitating molecular crosstalk during HNSCC pathogenesis

**DOI:** 10.1101/2020.09.22.303180

**Authors:** Sourabh Soni, Prince Anand, Mohit Kumar Swarnkar, Vikram Patial, Narendra V. Tirpude, Yogendra S. Padwad

## Abstract

**Background:** Transcriptome profiling has been pivotal in better comprehending the convoluted biology of tumors including head and neck squamous cell carcinoma (HNSCC). Recently, growing evidence has implicated the role of mitogen-activated protein kinase-activated protein kinase-2 (MAPKAPK2 or MK2) in many human diseases including tumors. MK2 has been recently reported as a critical regulator of HNSCC that functions *via* modulating the transcript turnover of crucial genes involved in its pathogenesis. Comprehensive MK2-centric transcriptomic analyses could help the scientific community to delve deeper into MK2-pathway driven mechanisms of tumor progression, but such studies have not yet been reported. Consequently, to delineate the biological relevance of MK2 and its intricate crosstalk in the tumor milieu, an extensive transcriptome analysis of HNSCC was conceptualized and effectuated with MK2 at the nexus.

**Methods:** In the current study, comprehensive next-generation sequencing-based transcriptome profiling was accomplished to ascertain global patterns of mRNA expression profiles in both *in vitro* and *in vivo* models of the HNSCC microenvironment. The findings of the RNA-sequencing analysis were cross-validated *via* robust validation using nCounter gene expression assays, immunohistochemistry, and real-time quantitative polymerase chain reaction (RT–qPCR).

**Results:** Transcriptomic characterization followed by annotation and differential gene expression analyses identified certain MK2-regulated candidate genes constitutively involved in regulating HNSCC pathogenesis, and the biological significance of these genes was established by pathway enrichment analysis. Additionally, advanced gene expression assays through the nCounter system in conjunction with immunohistochemical analysis validated the transcriptome profiling outcomes quite robustly. Furthermore, the results obtained from immunohistochemistry and transcript stability analysis indicated the crucial role of MK2 in the modulation of the expression pattern of these genes in HNSCC tumors and cells.

**Conclusions:** Conclusively, the findings have paved the way toward the identification of new effective tumor markers and potential molecular targets for HNSCC management. The results have accentuated the importance of certain differentially expressed MK2-regulated genes that are constitutively involved in HNSCC pathogenesis to potentially serve as putative candidates for future endeavors pertaining to diagnosis and therapeutic interventions for HNSCC.

## 1. Background

Multifaceted regulatory networks tend to connect genes within a myriad of cellular processes. A plethora of genes are involved in fundamental biological processes such as cell differentiation, growth, and programmed cell death, and their role in many diseases is presently known [1]. However, the apprehension of their roles at a global level is still incomplete. Gene transcription and regulatory networks in conjunction with new genome-wide approaches have garnered huge attention in the pretext of gene regulation. Nevertheless, post-transcriptional mechanisms such as transcript stability are also highly crucial and require intricate regulation *via* a multitude of intracellular signaling pathways [2, 3]. In particular, the modulation of transcript stability through phosphorylation-mediated regulation of RNA-binding proteins (RBPs) by mitogen-activated protein kinases (MAPKs) has been a topic of great interest [2, 3, 4, 5].

Head and neck squamous cell carcinoma (HNSCC) having an incidence rate of ∼600,000 cases yearly, is the seventh most common cancer worldwide and one of the most lethal cancers with an overall mortality rate of 40-50% [6,7]. HNSCCs are classified either histologically [8] or *via* the analysis of global transcription that employs etiology-specific profiles [9, 10]. However, when these parameters were used for patient clustering, specific differences were observed in the clinical behavior of patients as well as their response to therapy [11]. The survival rates of HNSCC patients have not improved much, hence, HNSCC has been rightly termed a malignant tumor with a low survival rate [12]. Consequently, augmented mechanistic insight into the molecular basis of HNSCC pathogenesis is urgently required to help in the early diagnosis and development of effective therapeutics aimed at improved clinical outcomes [13].

The role of differentially expressed genes (DEGs) and endogenous RNA networks in HNSCC is not fully deciphered. Past reports on genome and transcriptome studies in various human tumors have revealed aberrant regulatory programs, driver mutations, and disease subtypes [14]. The cancer genome study is a valuable tool for classification, diagnosis, and prognosis in HNSCC. There have been many past reports pertaining to genomic alterations in HNSCC [15, 16]. Recently, The Cancer Genome Atlas (TCGA) has led to a global analysis of major molecular changes, a comprehensive landscape of transcriptomic alterations and pathogenesis-linked signaling pathways in tumors, thus, contributing to the identification of novel prognostic biomarkers or specific anticancer molecular targets [17, 18]. However, there is still a need for extensive research insights to decipher the prognostic value attributed to these genomic alterations in tumors such as HNSCC. A variety of biomarkers such as MAPK phosphatase-1 (MKP-1), p16, p27, p53, tumor necrosis factor-α (TNF-α), and vascular endothelial growth factor (VEGF) have been shown to be linked with HNSCC [5, 19], but they have not been proven to be sufficient in accurately defining HNSCC pathogenesis. Single biomarkers have generally proven insufficient in the prediction of therapeutic response thereby necessitating research on combinatorial markers through high-resolution “omics” profiling [20]. Consequently, the identification of reliable molecular biomarkers associated with HNSCC using omics-based analyses is needed to develop novel potential diagnostic and therapeutic targets [21].

Recently, the mRNA regulatory networks involved in tumor progression have garnered huge research interest with recent reports showing the role of these intricate networks in tumorigenesis [22]. However, research endeavors in this area are quite limited, thereby pointing to a pertinent need for comprehensive analyses of mRNAs and regulatory networks and their involvement in tumorigenesis. Next-generation sequencing (NGS) has rapidly evolved as an important tool for epigenomic, genomic, and transcriptomic profiling of cancers. Technological advances in mining and deciphering vast transcriptomic data have enabled us to better comprehend the complexity of various tumors and have streamlined efforts to discover novel biomarkers and therapeutic targets aimed at tumor management [23]. In a recent study, our team elucidated the role of mitogen-activated protein kinase-activated protein kinase-2 (MAPKAPK2 or MK2) in HNSCC pathogenesis using clinical tissue samples, cell lines, and heterotopic xenograft mouse model [5]. MK2 was found to be critically important in regulating HNSCC *via* modulating the transcript stability of crucial pathogenesis-related genes. It was also established that MK2-knockdown attenuated tumor progression in a xenograft mouse model [5]. Thereupon, to delve deeper into the mechanistic role of MK2 and to decipher the molecular markers responsible for MK2-mediated changes in HNSCC pathogenesis, a comprehensive transcriptome profiling was performed and evaluated.

In the present study, the global mRNA expression profiles in HNSCC experimental model sets were evaluated using transcriptome analysis on the NovaSeq 6000 system (Illumina Inc., USA). The *in vitro* HNSCC cells, CAL27-MK2_WT_ (wild-type) and CAL27-MK2_KD_ (knockdown), cultured in normoxic or the tumor microenvironment mimicking hypoxic conditions comprised the first set. The *in vivo* heterotopic HNSCC xenograft bearing tumors from CAL27-MK2_WT_ and CAL27-MK2_KD_ cells in immunocompromised mice (as described previously [5] formed the second set. Comprehensive transcriptome analysis in the experimental models highlighted certain specific MK2-mediated DEGs and regulatory networks that play an integral role in HNSCC pathogenesis. Furthermore, specific gene expression assays on the nCounter system (NanoString Technologies, Inc., USA) were carried out to obtain a sensitive, highly multiplexed, and reliable detection of the defined mRNA targets based on the initial transcriptome profiling. The assays yielded highly precise and reproducible data that confirmed the transcriptome findings and yielded three MK2-regulated candidate genes (IGFBP2, MUC4, and PRKAR2B) intrinsically involved in HNSCC pathogenesis. Finally, cross-validation of the nCounter assay results in an *in vitro* setting, using immunohistochemistry (IHC) and mRNA transcript stability experiments through real time-quantitative polymerase chain reaction (RT-qPCR) was performed and it was found that these MK2-downstream genes showed dependence on MK2 for their expression and regulation in HNSCC tumors and cells (Figure 1). The findings corroborated recently published results, hence, ascertaining a crucial role of MK2 in HNSCC pathogenesis by means of transcript stability regulation [5]. These outcomes could potentially aid in the discovery of novel molecular markers for HNSCC management and diagnostic benefits.

**Figure 1:**
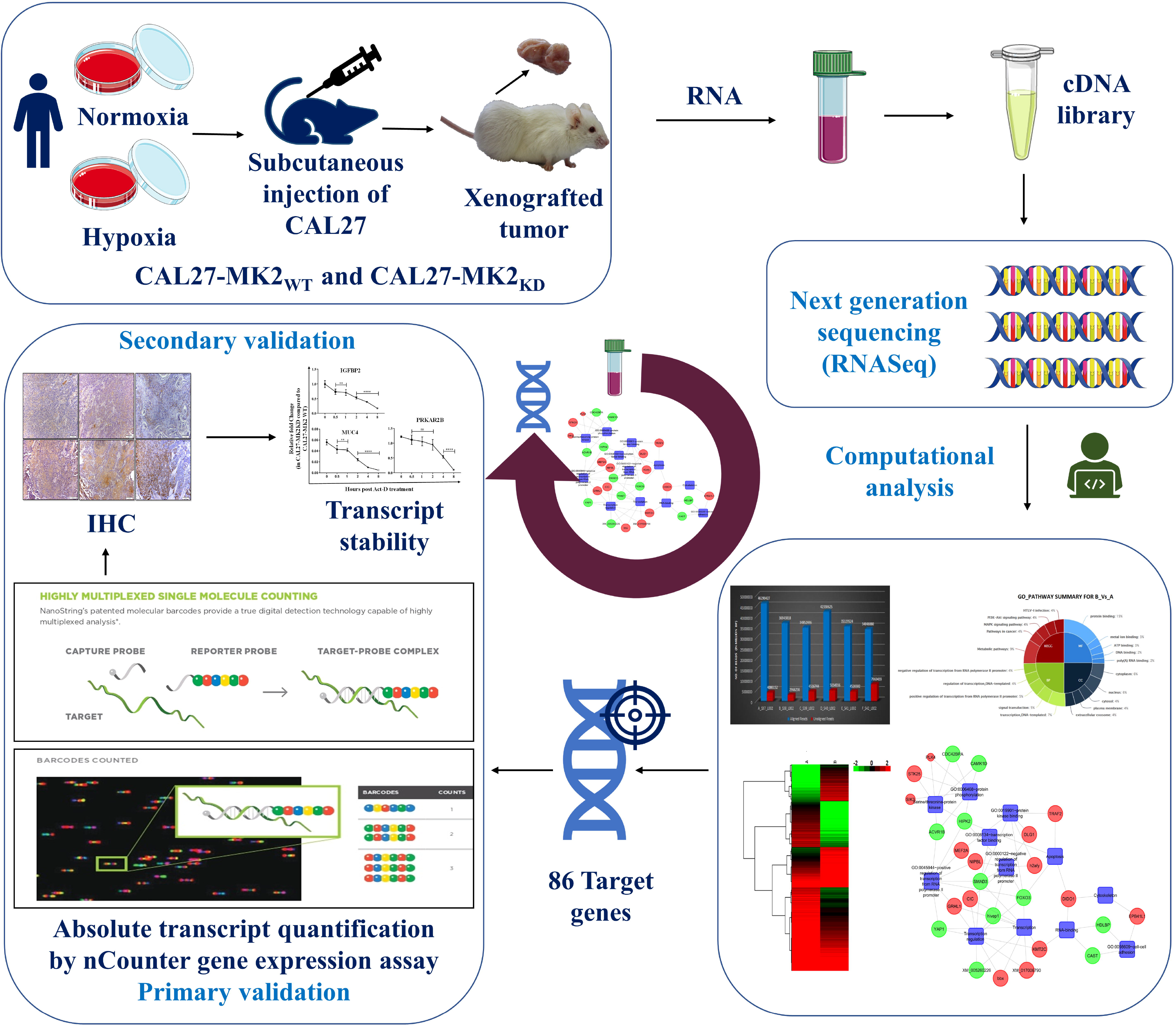
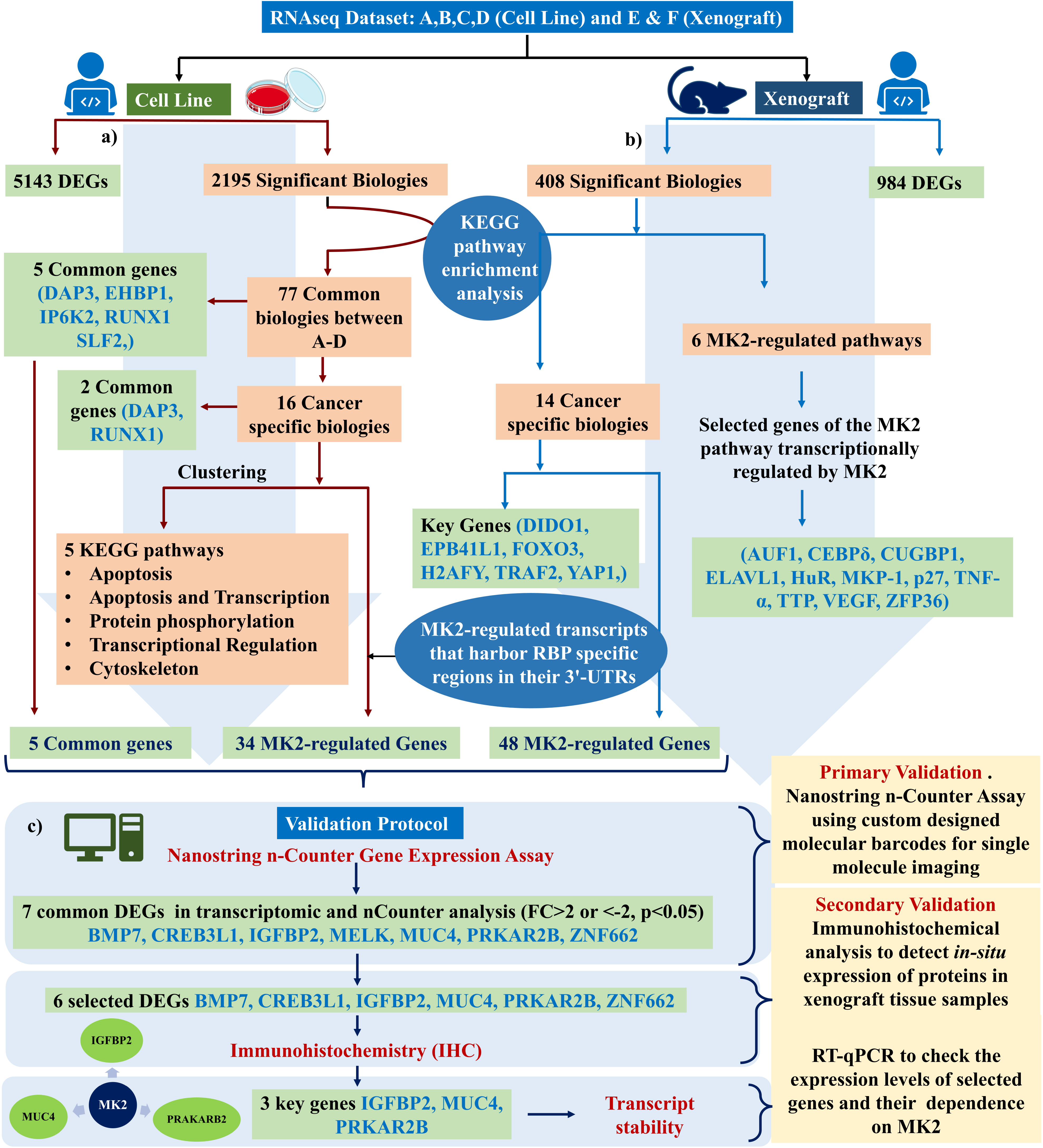
Schematic pictorial illustration of the analysis and validation workflow used for transcriptomic profiling and represents the key outcomes obtained with the work plan. Detailed analysis work plan and validation scheme indicating the two distinct sample types used for the experimental analysis and their output in terms of DEGs, significant biologies, important filtered out genes, and their validation. Analysis workflow of the dataset obtained from **a)** RNA-seq of cell line samples. **b)** RNA-Seq of tumor xenograft samples. This section represents the initial filtering of the transcriptome data for the identification of crucial biological processes and genes for all the experimental datasets. Additionally, expression levels and gene networks were checked for two categories of genes, first for the genes that are transcriptionally regulated by MK2 and second, for the genes having their mRNA stability controlled by MK2 through RBP-mediated regulation. Thereafter, MK2-regulated transcripts that harbor RBP specific regions in their 3’-UTRs were identified in all datasets, providing 34 genes in A to D comparisons and 48 in F vs E comparisons which were further analyzed by nCounter gene expression analysis. Finally, **c)** validation workflow of the MK2-regulated transcripts that harbor RBP specific regions in their 3’-UTRs filtered out from cell line and tumor samples, and 7 common DEGs in transcriptomic and nCounter gene expression assay were identified, 6 of them from the F vs E dataset were subjected to IHC validation and filtered 3 leads were subjected to transcript stability evaluation.

## 2. Materials and Methods

### 2.1 Cell Culture

*Homo sapiens* tongue squamous cell carcinoma cell line CAL27 (CRL-2095^™^, ATCC, USA) was grown in specific media supplemented with 10% fetal bovine serum and 1% antibiotic-antimycotic (Gibco, USA). The cells were cultured under normal conditions (37°C, 5% CO_2_ incubator with 95% humidity) and were free from any kind of contamination. Furthermore, MK2-specific short hairpin RNA-green fluorescent protein (shRNA-GFP) constructs were used to stably knockdown MK2 in cultured CAL27 cells to generate CAL27-MK2_KD_ cells as previously described [5]. For hypoxia exposure, the cultured cells were seeded into Petri plates and incubated in 0.5% O_2_ at 37°C in a hypoxia chamber for 48 hours (Bactrox, Shel-Lab, USA).

### 2.2 Xenograft mice model generation

To mimic the human tumor microenvironment, a biologically relevant heterotopic xenograft model of HNSCC was developed in non-obese diabetic/severe combined immunodeficient (NOD/SCID) mice. The immunocompromised mice were randomly assigned into control (CAL27-MK2_WT_) and experimental (CAL27-MK2_KD_) groups based on the specific cell type injected [5]. Briefly, for xenograft generation, 1×10^6^ cultured cells suspended in 100 μl of 1x phosphate buffered saline were injected subcutaneously into the right flanks of mice. Seven weeks post-graft inoculation, the mice were euthanized by CO_2_ asphyxiation; tumors were aseptically excised, weighed, and used for tissue embedding or RNA isolation.

### 2.3 RNA extraction and sample preparation for RNA-sequencing (RNA-seq)

CAL27-MK2_WT_ and CAL27-MK2_KD_ cells cultured in normoxia/hypoxia and tumors resected from the xenografted mice were employed for isolation of total cellular RNA using the RNeasy Mini kit (Qiagen, Germany) following the manufacturer’s recommended protocol (sample details are provided in Table 1). Consequently, the qualitative and quantitative assessment of all the RNA samples was performed using a NanoDrop 2000C spectrophotometer (Thermo Fisher Scientific, USA) and Bioanalyzer (Agilent 2100, Agilent Technologies, USA) (Figure S1). The RNA integrity number (RIN) value >5 was used as an exclusion criterion for this study. RNA samples having RIN>5 was used for cDNA library preparation. For each sample, RNA was isolated from at least three biological replicates for library construction and further experimentation.

### 2.4 cDNA library preparation and sequencing

Total RNA (5 µg) from each sample was used to isolate poly-A mRNA followed by preparation of cDNA library using the TruSeq mRNA sample preparation kit v2 (Illumina Inc.). Each sample was tagged with a unique TruSeq index tag to prepare multiplexed libraries. Six paired-end adapters with unique six base index sequences, permitting accurate differentiation among samples, were used for the library preparation. The quantification of prepared libraries was performed on a Qubit fluorometer using a Qubit dsDNA BR assay kit (Life Technologies, USA), while the size and purity of the libraries were examined on a Bioanalyzer DNA 1000 series II chip (Agilent Technologies). The flow chart of the sequential steps involved in the TruSeq library preparation is given in Figure S2. The libraries (4 from the cell line model and 2 from the animal model) had an average insert size of 210 base pairs (bp) and were pooled by taking 10 µl from each library. The final pool was loaded in one lane of an S2 flow cell using the NovaSeq XP protocol (Illumina Inc.) (Figure S3) [24]. Cluster amplification and generation of sequencing data were performed on the NovaSeq 6000 system (Illumina Inc.) using 2×100 paired-end cycles. Raw data quality control was accomplished using the NGSQC tool kit v2.3 with default parameters [25].

### 2.5 Reference-based assembly and homology search

The raw FASTQ files with low-quality reads of sequencing data were filtered to obtain high-quality filtered data that were aligned to the reference genome (genome reference consortium human build 38 patch release 12, GRCh38.p12). The Kallisto pipeline was used for alignment and identification of transcript coding regions followed by quantitation and annotation using default parameters [26][27]. Furthermore, the removal of multi-mapped reads was performed, and the filtered data were finally converted to read counts for annotated genes. Figure 2a is a flowchart representation of the various steps involved in the sequencing data analyses.

**Figure 2:**
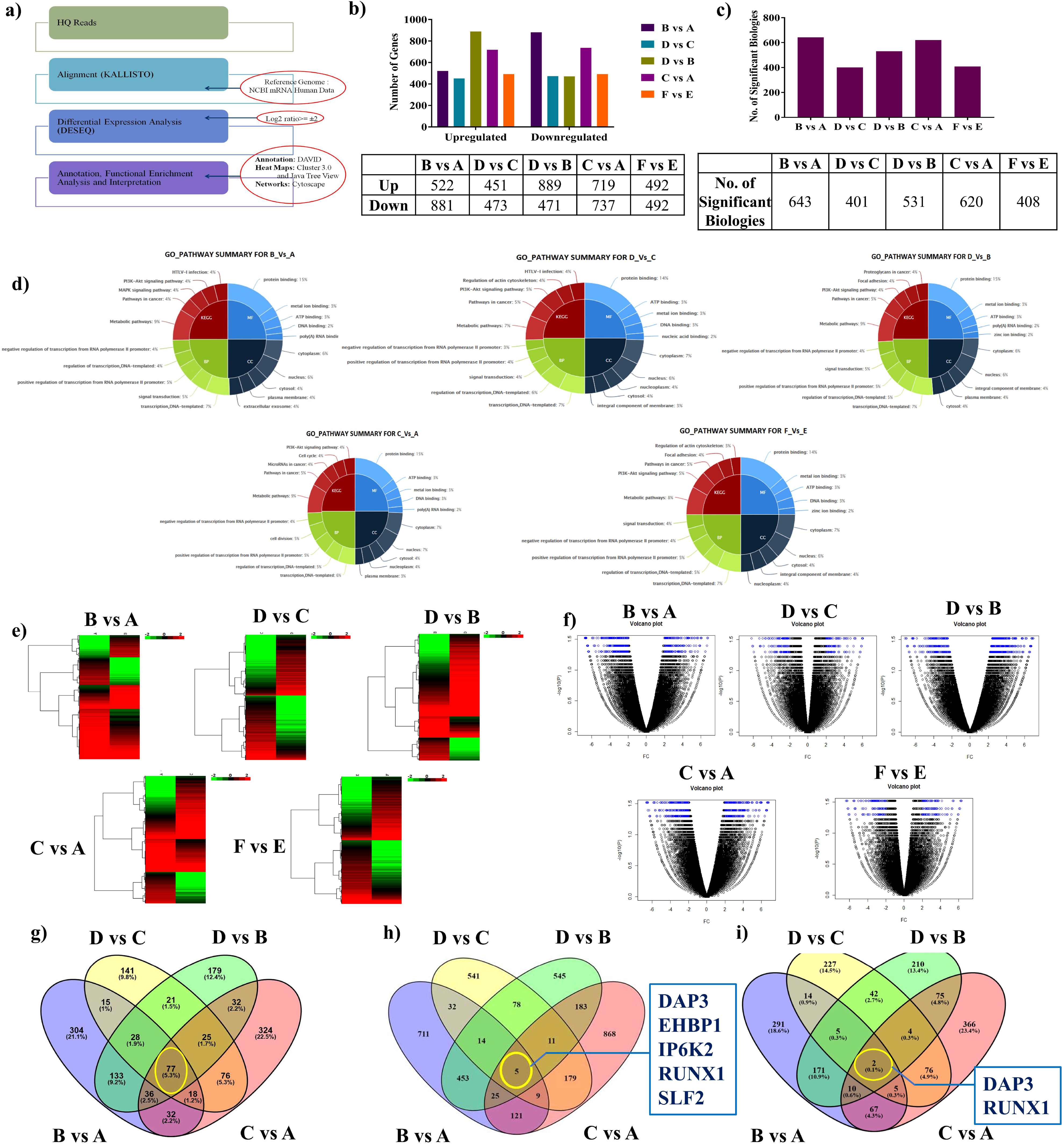
**a)** Workflow of sequencing analysis showing sequential steps and various tools and pipelines employed for transcriptome profiling. **b), c)** A comparative bar diagram representation of the number of (b) DEGs; and (c) Significant biologies/Biological processes, present in various datasets in the transcriptome profiling study**. d)** Pie chart representation of the top five GO terms and pathway summary based on all the DEGs in the various datasets showing approximately 5% of the total DEGs belonging to the pathways involved in cancer progression. **e)** Non-hierarchical heatmap representation depicting the expression profile and variation in average log2 fold change among DEGs in various datasets in the transcriptome profiling study. The color bar represents the expression values with green representing the lowest (downregulation) and red representing the highest (upregulation) expression levels. The various datasets used for expression profiling are labeled on the top. **f)** Volcano plot representation of the complete transcript list according to their average log2 fold change and p-values for various datasets in the transcriptome profiling study with differential transcripts highlighted in blue. The plot displays DEGs along the dimensions of biological significance (average log2 fold change) and statistical significance (p). Genes with an absolute log2 fold change>2 and a p-value<0.05 were considered as DEGs. **g)** Venn diagram representation created using Venny 2.1.0 showing the 77 common elements in the transcriptome profiling of the *in vitro* HNSCC cell line model (A-D datasets). **h)** Venn diagram representation created using Venny 2.1.0 showing the five common genes in the 77 common elements in the transcriptome profiling of the *in vitro* HNSCC cell line model (A-D datasets). **i)** Venn diagram representation created using Venny 2.1.0 showing the two common genes in the 16 common elements in the transcriptome profiling of the *in vitro* HNSCC cell line model (A-D datasets).

### 2.6 Annotation, differential gene expression, and pathway analyses

Expression of the transcripts in the samples was analyzed based on their fragments per kilobase of transcript per million mapped (FPKM) values [26]. Transcripts were given a score for their expression by the Cufflinks-based maximum likelihood method and values with FPKM≥0.1 were considered significant for downstream analysis. Although FPKM≥0.1 cut-off indicates a low level of transcript expression, this value was essentially used to attain a high enough threshold for the number of transcripts in the analyzed datasets considering the downstream filter-specific analyses performed in this study. Transcripts uniquely expressed in each sample were considered specific and were analyzed separately. The false discovery rate (FDR) was employed to correct the statistical significance of the p-values for multiple tests. DEGs in the analyzed datasets were identified *via* the DESeq analysis pipeline [27] using a fold change (FC) threshold of absolute log2 FC≥2 and a statistically significant Student’s t-test p-value threshold adjusted for FDR<0.001. Consequently, transcripts with FC<-2 were considered downregulated while those with FC>2 were considered upregulated. Statistically, significant enriched functional classes with a p-value adjusted for FDR>0.05, derived using the hypergeometric distribution test corresponding to the DEGs, were determined using Student’s t-test with the Benjamini Hochberg FDR test.

Unsupervised hierarchical clustering of DEGs was performed using Cluster 3.0 and visualized using Java TreeView [28, 29]. Gene ontology (GO) and pathways that harbor expressed transcripts were identified using the DAVID functional annotation tool (http://david.abcc.ncifcrf.gov/home.jsp) [28, 29, 30] (Figure 2a). For the DEGs, heat maps and volcano plots were generated using the ‘gplots’ and ‘heat map’ packages. GO and Kyoto Encyclopedia of Genes and Genomes (KEGG) pathway analyses were performed for the assembled transcripts with reference to the UniProt database. Data of the total DEGs (upregulated and downregulated) were explored further using Cytoscape v3.5.0 (http://www.cytoscape.org/) to better understand the gene regulatory networks and for mapping of the results [31]. In figures depicting the gene regulatory networks, the gene nodes/circles are sized according to their p-values and colored according to their FC where red shows upregulation, green depicts downregulation, and yellow displays baseline expression; processes are shown in rectangular boxes and colored in blue.

### 2.7 The nCounter gene expression assays for primary validation

To validate the leads obtained from the transcriptomic profiling, custom-designed molecular barcodes (NanoString Technologies, Inc.) were utilized for single-molecule imaging, thereby making it possible to detect and count hundreds of different transcripts in one reaction (Figure 1 and S4). RNA quality was assessed using the Agilent 2100 Bioanalyzer (Agilent Technologies) (Figure 1 and S1). Gene expression was analyzed on the nCounter system (NanoString Technologies, Inc.) following the manufacturer’s recommendations. Briefly, the custom synthesized probes were hybridized overnight to the target RNA followed by washing away of the excess probes, immobilization of the CodeSet/RNA complexes in the nCounter cartridge, and finally data collection on the nCounter system (Figure 1 and S4). The gene expression levels were measured in triplicate for total RNA from the cell line and xenografted tumor samples, normalized to the four housekeeping genes (HKGs), and analyzed using nSolver software (NanoString Technologies, Inc.). Each nCounter assay contained synthetic spike-in controls in the preparatory mix to allow correction of the sample-to-sample variation arising due to common experimental errors such as differences in the amount of input transcripts or reagents [32]. The counts were normalized to the positive controls and averaged for the samples of each mRNA type. Normalization involved spiked-in positive and negative control probes for background correction in addition to the 4 HKGs. Data analyses were performed on the nSolver 3.0 analysis software (NanoString Technologies, Inc.).

### 2.8 Immunohistochemical analysis for secondary validation

The levels of expression and activation status of specific proteins were analyzed using IHC in the tumor sections from the *in vivo* xenograft model to validate the findings of the transcriptomic and the nCounter gene expression analysis. The animal study was approved by the Institutional Animal Ethics Committee (IAEC) of CSIR-IHBT, Palampur, India (Approval No. IAEC/IHBT-3/Mar 2017). IHC was performed according to the previously reported protocol [5]. Briefly, 5 μm thin sections fixed on poly-l-lysine coated slides were deparaffinized and rehydrated. Antigen retrieval was performed using sodium citrate buffer (pH 6.0) followed by quenching of endogenous peroxidases using BLOXALL^TM^ blocking solution (Vector Laboratories, Inc., USA). Furthermore, incubation of the sections with 2.5% normal horse serum blocked the exposed sites. Sections were then incubated with appropriately diluted specific primary antibody (Table ST1) overnight at 4°C followed by horseradish peroxidase-conjugated secondary antibody for 1 hour at room temperature. The rinsed sections were then incubated with 3,3’-diaminobenzidine substrate and Mayer’s hematoxylin served as a counterstain. Five field views were obtained from each section for the designated antibodies and used for quantitative analysis of protein expression.

### 2.9 Gene expression analysis and MK2-regulated transcript stability

To determine the role of MK2 in regulating transcript stability, the expression levels of selected genes and the stability of their transcripts were assessed in the presence/absence of MK2 using RT-qPCR in CAL27 cells. shRNA constructs were used to generate CAL27-MK2_KD_ cells as previously reported [5]. MK2-knockdown was confirmed using western blotting (WB), following which CAL27-MK2_WT_ and CAL27-MK2_KD_ cells were treated with 1 µM actinomycin-D (Act-D) to inhibit transcription. Total RNA was isolated at 6 time points (0, 0.5, 1, 2, 4, and 8 hours) using the RNeasy Mini kit (Qiagen) according to the manufacturer’s recommended protocol. The purity of the isolated RNA was determined using a NanoDrop 2000C spectrophotometer (Thermo Fisher Scientific) and the 260/280 ratios were found to be between 1.9 and 2.1. Further RT-qPCR was performed using a Verso 1-step RT-qPCR kit (Invitrogen, USA) as previously described [5]. GAPDH was used as an internal gene control and the difference in cycle threshold (Ct) was calculated following the 2^-ΔΔCt^ method. The relative fold change was calculated by comparing the CAL27-MK2_KD_ (designated as shMix group, cells were treated with a combination of an equal amount of MK2 targeting shRNA complexes 1, 2, 3, and 4) at the 6 mentioned time points with CAL27-MK2_WT_ (designated as mock or scramble control group, cells were treated with scrambled shRNA). shRNA constructs and vector maps have been described previously [5].

### 2.10 Statistical analysis and quantification

All the experiments were conducted at least in triplicates unless mentioned otherwise. IHC staining intensity was observed and analyzed by an expert pathologist in a single-blinded fashion using a BX53 bright field microscope (Olympus Corporation, Japan). Quantification of protein expression for IHC and WB was performed using ImageJ software 1.8.0 (https://imagej.nih.gov/ij/). The images were RGB stacked, and the color threshold was adjusted according to the expression of specific proteins in the tissue section. For quantification, the expression was presented as a composite score of the percent area of the total tissues using ImageJ software. The statistical/imaging parameters used for the various transcriptomic analyses have been detailed with their explanations in their respective sections in the manuscript. GraphPad Prism 7.0 software (GraphPad Software, Inc., USA) was used for the data analysis of IHC and RT-qPCR quantified datasets.

## 3. Results

### 3.1 Qualitative assessment of the generated cDNA library followed by filtering and assembly of reads depicted optimum alignment

The quality assessment of the isolated total-RNA from the appropriate cells/tissues, as well as the cDNA library generated, was performed using a Bioanalyzer (Agilent 2100, Agilent Technologies). It was found that the RIN values of all the RNA samples and the cDNA library were >5 suggesting that they were of suitable quality for use in downstream experiments (Figure S1 and S3).

Deep sequencing of RNA obtained from the NovaSeq 6000 platform (Illumina Inc.) resulted in 349 million raw reads (∼58.2 million raw reads per sample) with an average insert size of 210 bp. Figure S5 and S6 summarizes the quality check (QC) results of the sequencing experiment. The raw FASTQ sequences were filtered using the NGSQC tool kit to obtain high-quality (HQ) reads based on the predefined parameters, generating 258 million filtered HQ reads (∼43.1 million HQ reads per sample). This amounted to 74.1% of the total raw reads, implying that the obtained data were of good quality. A total of 25.7 gigabases of data were generated which enumerates the enormity and wide complexity of the human genome.

The HQ reads obtained were further examined in the downstream analyses and mapped over the human reference genome. The alignment performed by employing the Kallisto pipeline was optimum with 88.7% of the HQ reads on average being mapped to the human reference genome (Figure 2a and S6).

### 3.2 Annotation and differential gene expression analyses revealed candidate genes

The present study was conducted using two distinct experimental datasets as mentioned in Figure 1 and Table 1. Using the criteria of FPKM≥0.1, 62791 expressing transcripts on an average were identified per sample, which further represented thousands of genes. A-D datasets depict the *in vitro* HNSCC cell line samples while F vs E illustrates the *in vivo* xenograft dataset. As detailed in Figure 1 and Table 1, analyses of MK2_KD_ vs MK2_WT_ in normoxia and the tumor core emulating the hypoxic niche were performed to obtain a comprehensive picture of the changes in the global gene expression pattern. The differential gene expression in the analyzed datasets was evaluated and a large pool of the DEGs, precisely 1403 in B vs A, 924 in D vs C, 1360 in D vs B, 1456 in C vs A, and 984 in F vs E were found as assessed by the predefined cutoff values. Figure 2b represents the total number of upregulated (FC>2) and downregulated (FC<-2) genes among all the DEGs in the analyzed datasets.

### 3.3 Pathway enrichment analysis revealed the biological significance of the findings

The multitude of DEGs in the transcriptome profiling datasets was implicated in hundreds of significant biologies/biological processes as summarized in Figure 2c. To gain further insight into the biological significance of the variations in gene expression and to attain a global picture of the molecular pathways possibly contributing to HNSCC pathogenesis, pathway enrichment analyses were performed using the KEGG database. The resultant integrated network analysis revealed the top biological processes enriched in the analyzed datasets. The top GO pathway analyses for the DEGs are depicted in Figure 2d. Notably, a significant percentage (∼5%) of the total DEGs in the analyzed datasets belonged to the pathways involved in cancer progression (Figure 2d). The DEGs showing significant changes among various groups were then selected, followed by the generation of heat maps to assess the clustering of gene expression profiles among the analyzed datasets (Figure 2e). Figure 2f further depicts the distribution of all the transcripts on the two dimensions of -log(P) and FC by way of volcano plots with differentially expressed transcripts highlighted in blue.

In the A-D datasets, the DEGs belonged to a multitude of biological processes thereby limiting the information that could be harnessed. Henceforth, to filter down the data to attain meaningful outcomes and fulfill the aim of extracting valuable leads, the 77 elements/biological processes that were common in A-D datasets were selected using Venny 2.1.0 (http://bioinfogp.cnb.csic.es/tools/) (Figure 2g). These processes belonged to the various categories listed in Table ST2. Based on this knowledge, the common genes in these 77 biological processes were further filtered down using the same approach (Figure 2h). Consequently, 5 genes common in the A-D datasets were obtained namely death-associated protein 3 (DAP3), EH domain binding protein 1 (EHBP1), inositol hexakisphosphate kinase 2 (IP6K2), runt related transcription factor 1 (RUNX1), and SMC5-SMC6 complex localization factor 2 (SLF2) as listed in Table ST3. It is worth mentioning here that these MK2-regulated genes portray essential roles in HNSCC pathogenesis [33, 34, 35, 36]. Hence, further exploratory investigations of these putative molecular targets are deemed essential for elucidating their relevance in HNSCC management.

### 3.4 Gene regulatory networks and pathways depicted the significance of the candidate genes in HNSCC pathogenesis

Gene regulatory networks for the 5 common genes in A-D datasets furnished a detailed overview of the various inter-network connections and the biological processes affected by them. Collectively, the results showed that DAP3 plays an intrinsic role in apoptosis and poly(A)-RNA binding while IP6K2 plays a role in ATP, nucleotide, and protein binding as confirmed in past reports [19, 32]. These genes showed differential regulation in *in-vitro* dataset (B vs A) hence, clearly signifying that MK2-knockdown, as well as the hypoxic tumor microenvironment, affects the genes and pathways *via* differential regulation, pointing to a central role of MK2 in transcriptional regulation of HNSCC. Interestingly, the *in-vitro* results published by our group recently [5] also pointed to the role of MK2 in modulating the transcript stability of MK2-regulated genes. Similarly, the xenograft dataset (F vs E) potentiated this finding (Figure 3a, 3b).

**Figure 3:**
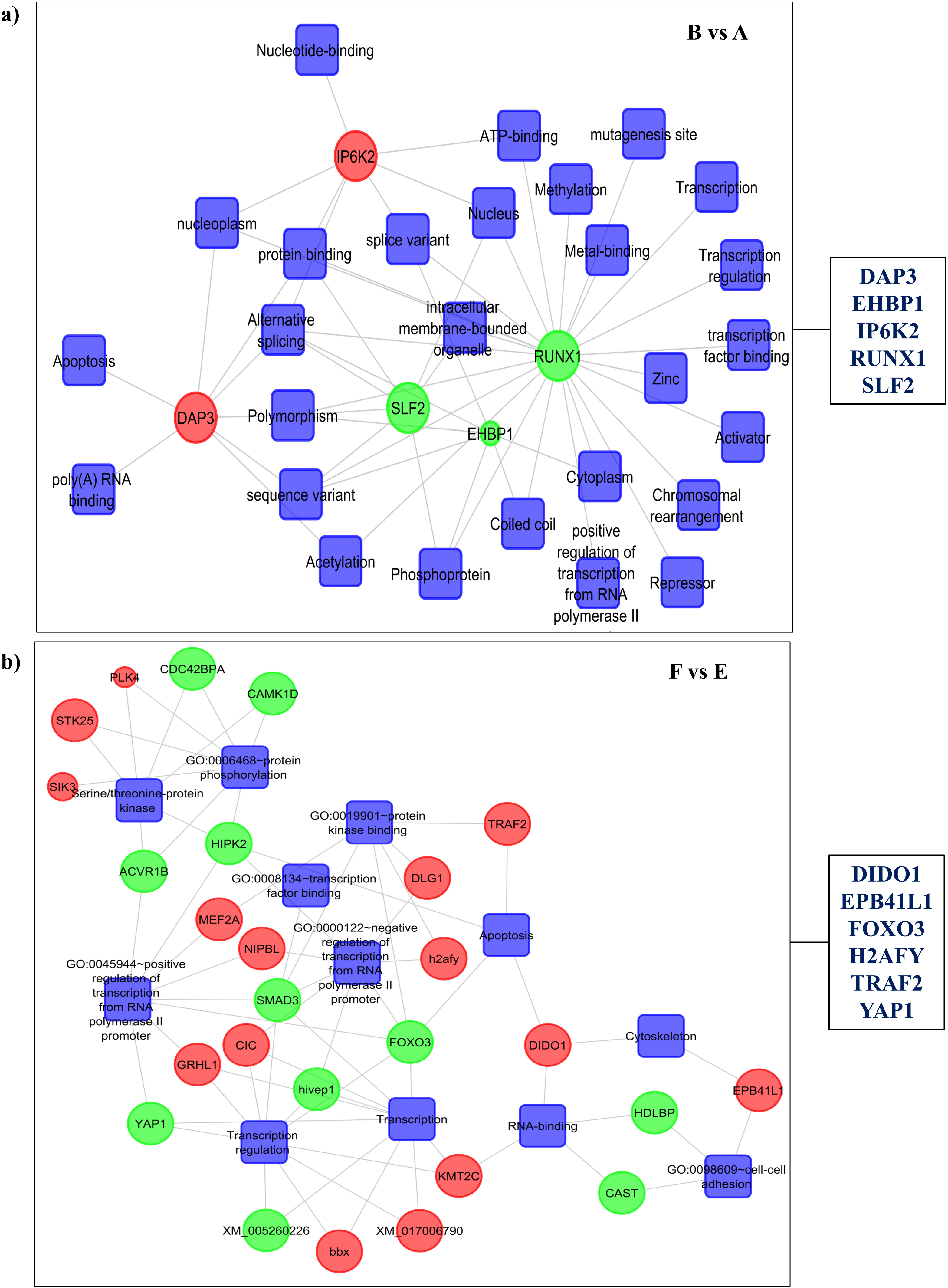
Gene regulatory network depicting a detailed overview of the various interconnections and the significantly enriched biological processes affected by the 5 common genes (DAP3, EHBP1, IP6K2, RUNX1 SLF2). Network represented in 77 common elements in the datasets where MK2 knockdown is present; **a)** B vs A comparison of the transcriptome profiling of the *in vitro* HNSCC cell line model. **b)** Gene regulatory network depicting a detailed overview of the various interconnections and the significantly enriched biological processes affected by the DEGs in 14 cancer-specific biological processes in the transcriptome profiling of the *in vivo* heterotopic HNSCC xenograft dataset (F vs E comparison). The gene nodes are sized according to their p-values and colored according to their average log2 fold change, where red shows upregulation while green shows downregulation; processes are shown in rectangular boxes and colored in blue.

Furthermore, to retrieve the information pertaining to the MK2-regulated candidate genes intrinsic to HNSCC pathogenesis, the data were narrowed down to include only processes that were involved in tumor progression for further analysis. This filtering down the dataset to 16 cancer-specific biological processes that were common in the A-D datasets (listed in Table ST4). In these 16 processes only 2 genes, DAP3 and RUNX1, were found to be common based on the analysis performed using Venny 2.1.0 (Figure 2i and Table ST5). Collectively, these findings clearly indicated that DAP3 and RUNX1 were differentially expressed in the cell line-based datasets analyzed, hence, potentiating their intrinsic involvement in HNSCC pathogenesis and warranting further investigation. Notably, the 16 processes were found to be clustered in 5 major biological pathways (Figure S7) including apoptosis and transcription, thereby, clearly showing that MK2 portrays an intrinsic role in the hypoxic tumor microenvironment by regulating these processes, hence, substantiating the latest *in vitro* findings from our group [5].

Similarly, on comparing the mice bearing CAL27-MK2_KD_ tumors with those bearing CAL27-MK2_WT_ tumors (F vs E xenograft dataset), it was found that the DEGs were clustered in 14 biological processes relevant to tumor pathogenesis as listed in Table ST6. The genes involved in these biological processes were assessed and Figure 3b shows the gene regulatory network. This analysis provided certain MK2-regulated candidate genes, such as TRAF2 [34] (apoptosis); EPB41L1 [35] (cytoskeleton); FOXO3, H2AFY and YAP1 (transcriptional regulation) [36] [37] [38]; and DIDO1 (RNA binding) [39] which are involved in key cellular processes in the xenograft model (Figure 3b). These findings can be explored further to decipher the putative role of the potential candidate genes in HNSCC pathogenesis with an aim to define the probable therapeutic targets for HNSCC management.

### 3.5 MK2 is the master of regulatory networks and functions by modulating transcript stability

The prime objective of this study was to gather comprehensive information regarding the various MK2-regulated DEGs and pathways that render essential roles in HNSCC pathogenesis in normoxic and tumor core mimicking hypoxic conditions. Thereupon, keeping MK2 at the nexus of further analyses, this study focused on elucidating the regulation of the MK2 pathway and its downstream targets in the analyzed datasets. MK2 was found to be involved in the regulation of major biological pathways as shown in Figure S7 and S8. Recent findings by our group have asserted that MK2 controls the transcript stability of critical genes involved in HNSCC pathogenesis *via* RBP-mediated regulation [5]. Hence, we further analyzed the transcriptomic data to decipher the role of MK2 in the regulation of mRNA stability. Interestingly, MK2 was found to accomplish this task through RBP-mediated regulation with HuR (ELAVL1) and TTP (ZFP36) playing intrinsic roles (Figure 4a), thus, clearly affirming the hypothesis and corroborating previous findings [5]. The levels of expression of these RBPs and hence regulation varied in the analyzed datasets, thereby, clearly suggesting that the tumor microenvironment, in association with the presence/absence of MK2, plays an important role in HNSCC pathogenesis.

**Figure 4:**
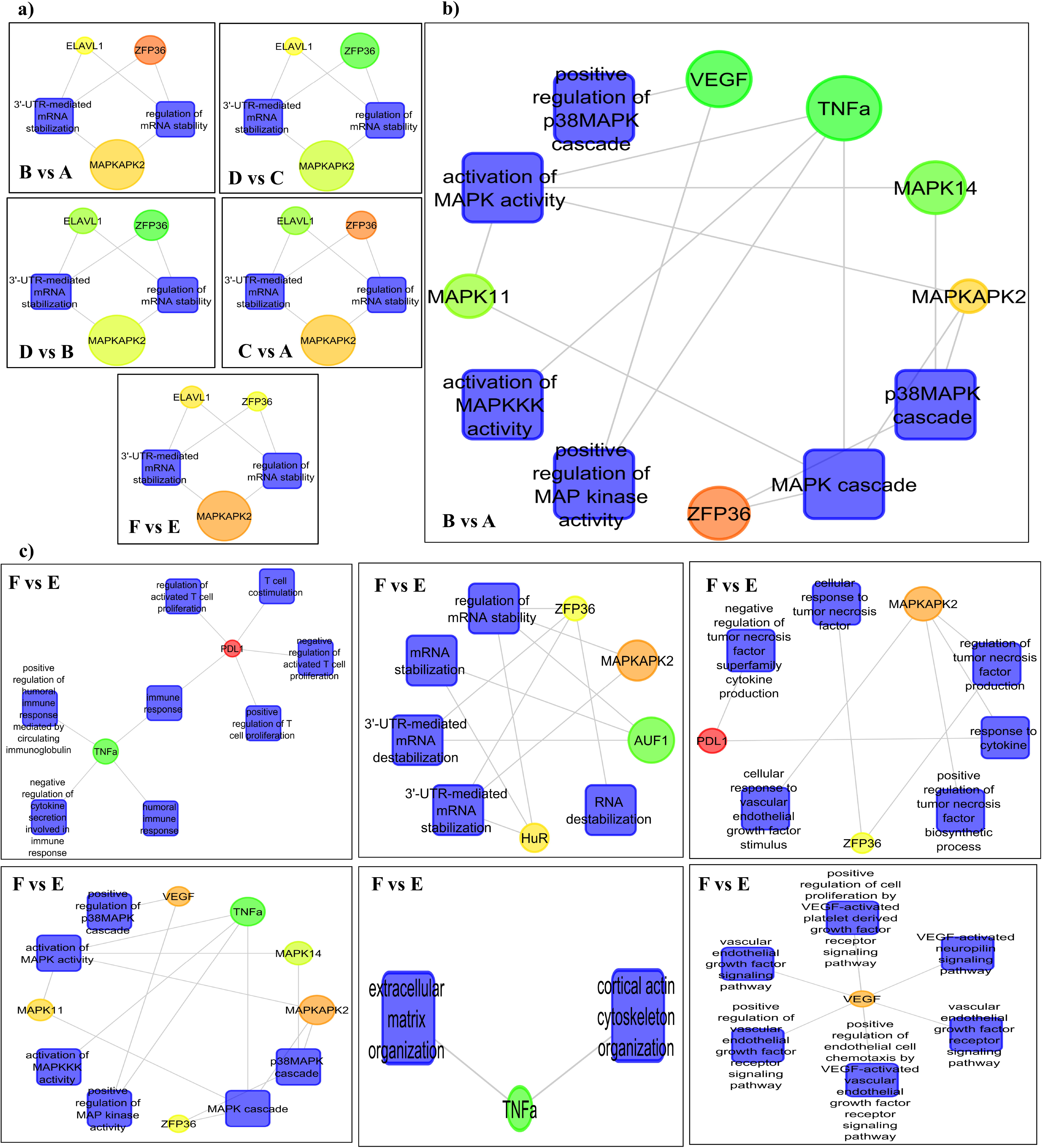
**a)** Representation of a gene regulatory network depicting the role of MK2 in the regulation of mRNA stability in the various datasets. The figure clearly demonstrates that MK2 regulates transcript stability *via* RBP-mediated regulation with HuR (ELAVL1) and TTP (ZFP36) playing an integral part. **b), c)** Gene regulatory network showing MAPK signaling cluster of the selected MK2 pathway genes (p38, MK2, AUF1, TTP, CUGBP1, CEBPδ, HuR, MKP-1, p27, TNF-α, and VEGF) in the transcriptome profiling data of the (b) *in vitro* HNSCC cell line dataset (B vs A, normoxic microenvironment) indicating VEGF and TNF-α downregulation. (c) In F vs E comparison, these genes which portray essential roles in the MK2 pathway are playing intrinsic roles in certain cellular pathways that are clustered into 6 major biological processes such as immune response, regulation of mRNA stability, regulations of cytokines, MAPK signaling, cell migration, and signaling pathways. The gene nodes are sized according to their p-values and colored according to their average log2 fold change, where red shows upregulation while green shows downregulation and yellow indicates baseline expression; processes are shown in rectangular boxes and colored in blue.

Next, to attain a clear understanding of what was happening at the transcriptional level, the analysis was narrowed down by selecting specific genes of the MK2 pathway. The genes were selected by literature mining of past MK2-centric studies and included those that were analyzed in our recent study [5]. Interestingly, the genes selected for this analysis, viz. AUF1, CEBPδ, CUGBP1, HuR, MK2, MKP-1, p27, p38, TNF-α, TTP, and VEGF, have been previously shown to be involved in several key cellular processes [reviewed in [4]. Elucidation of the MAPK signaling cluster in detail indicated that in the background of MK2-knockdown in normoxia (B vs A dataset), TNF-α and VEGF tended to show downregulation (Figure 4b), which is in complete consonance with previously published results. The analyzed genes were clustered into 6 major biological processes as shown for (F vs E xenograft dataset) in Figure 4c and Figure S8. Collectively, the results of the transcriptomic analysis corroborated very well with previous findings, thereby, robustly verifying the hypothesis that MK2 is the master regulator of the transcript stability of genes critical to HNSCC pathogenesis.

### 3.6 3’-untranslated region (3’-UTR)-based filtering furnished information regarding important MK2-regulated downstream target genes

In the present study, we focused on the role of MK2 and MK2-regulated genes in HNSCC pathogenesis. It is well known that MK2 can potentially regulate the transcript stability of only those downstream targets that possess binding regions for RBPs in their 3’-UTRs [40]. Hence, the transcriptomic analysis was narrowed down to only those DEGs that harbored RBP-binding regions in their 3’-UTRs (Figure 5), an approach that has lately been the cornerstone of many ‘omics’ studies [41, 42, 43]. The DEGs were filtered based on the presence of adenylate-uridylate-rich elements (ARE)-regions in their 3’-UTRs where RBPs can potentially bind and modulate their function possibly *via* MK2-mediated regulation. To accomplish this task, the 3’-UTR regions of all the DEGs were fetched using Ensembl (http://www.ensembl.org/) [44]. Next, the domain sequences of RBPs were assessed using the catalog of inferred sequence binding preferences of the RBPs (CISBP-RNA) database [45]. Last, the transcripts that harbor RBP-specific regions in their 3’-UTRs were filtered out using the RBPmap v1.1 web tool (http://rbpmap.technion.ac.il/) (Figure 5a) [46]. Once the probable MK2-downstream targets were identified (based on the aforementioned approach), the data were reassessed focusing on the 16 previously selected cancer-specific pathways in the *in vitro* HNSCC cell line model (A-D datasets) (Table ST4). The top 2 upregulated and downregulated genes in this dataset were evaluated following 3’-UTR filtering which yielded 34 putative MK2-regulated genes as listed in Table ST7 (Figure 5b). Similarly, the topmost upregulated and downregulated genes were analyzed in all the cancer-specific pathways for the *in vivo* heterotopic HNSCC xenograft dataset (F vs E dataset) which provided 48 MK2-regulated genes that are listed in Table ST8 (Figure 5c). Collectively, this 3’-UTR-specific filtering of the transcriptomic data brought into the limelight possible MK2-downstream target genes that could be integral in HNSCC pathogenesis. Furthermore, to cross-validate the findings of the transcriptomic profiling, these candidate genes along with the 5 common genes in the 77 common elements in A-D datasets (listed in Table ST3) were used for further *in vitro* validation. H2AFY was common in the transcriptomic analysis for both the cell line and xenograft analysis (Table ST7 and ST8). Resultingly, a total of 86 genes (Table ST9) (34 MK2-regulated genes and 5 common genes for the A-D datasets, 48 MK2-regulated genes in the F vs E dataset, and the common gene, H2AFY, were counted once) were selected for further experimental validation (Figure 1).

**Figure 5:**
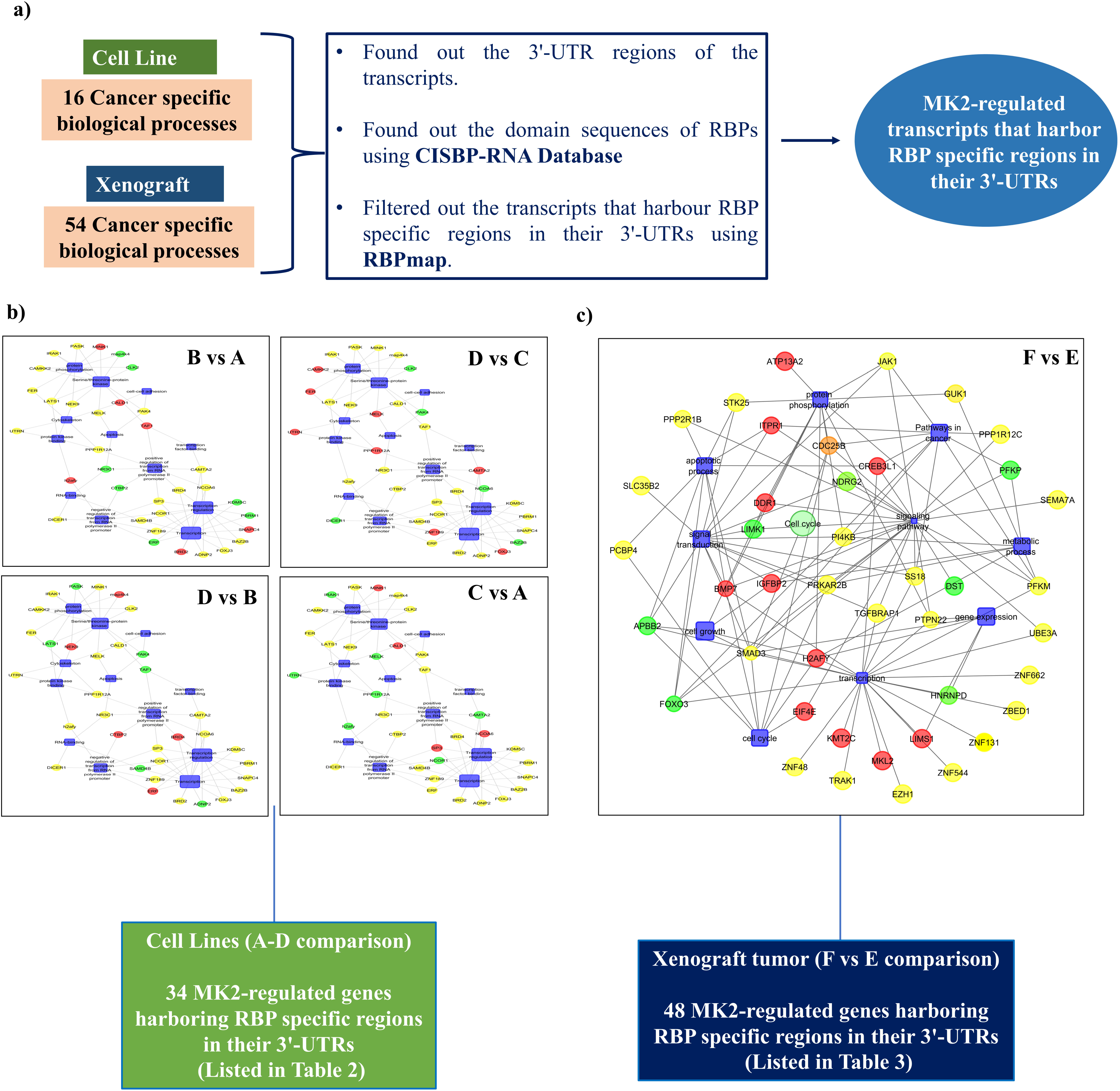
3’-UTR based filtering of cancer-specific biological processes from a cell line (16 processes) and xenograft tumor (54 processes) datasets to identify MK2-regulated transcripts that harbor RBP specific regions in their 3’-UTRs. **a)** Schematic representation of the applied workplan. **b)** and **c)** Regulatory network of MK2-regulated genes harboring RBP specific regions in their 3’-UTRs; b) total 34 in the B vs A, D vs C, D vs B and C vs A datasets, and c) total 48 in the F vs E datasets. The gene nodes are sized according to their p-values and colored according to their average log2 fold change, where red shows upregulation while green shows downregulation and yellow indicates baseline expression; processes are shown in rectangular boxes and colored in blue.

### 3.7 Highly efficient and precise detection of gene expression *via* nCounter gene expression assays potentiated transcriptomic outcomes

Routinely, the findings of transcriptomic analyses are generally validated in an *in vitro* setting *via* gene expression analysis (employing RT-qPCR) using the same RNA sample to maintain homogeneity. In lieu of the high-throughput nature of the validation in this study, RT-qPCR analysis could have been very tedious and prone to numerous errors. Hence, as a viable and more pragmatic alternative, the latest and highly precise gene expression assay-based nCounter system approach (NanoString Technologies, Inc.) was employed to validate the outcomes of the transcriptome analysis in this study. To accomplish this, 90 specific custom-designed molecular probes corresponding to the selected MK2-regulated candidate genes were procured to aid in the imaging and fast detection of multiple transcripts (90 in this study) in a single reaction with a high-fidelity rate (NanoString Technologies, Inc.). The gene set comprised 86 selected genes from the transcriptomic profiling as well as 4 HKGs (listed in Table ST9 and ST10, respectively). 4 commonly used reference genes, viz. ABCF1, GAPDH, POLR2A, and RPL19, were selected based on an extensive literature survey and because of their baseline expression in both HNSCC as well as in MK2-knockdown conditions [47, 48].

The assay was performed using the standard procedure as highlighted in Figure S4 and S9 and detailed in the methods section. Briefly, the custom synthesized probes were hybridized to the target RNA samples followed by washing off the excess probes. Further, immobilization of the probe/RNA complexes on the nCounter cartridge was performed, samples were run on the nCounter instrument followed by data retrieval (NanoString Technologies, Inc.). Some of these genes were differentially expressed in the analyzed datasets as indicated by the statistical analysis (p<0.05). Next, to make sense of the biological significance of changes in gene expression in the nCounter data, KEGG pathway enrichment analysis was performed that revealed the top 5 biological processes (Figure 6a and 6b). Furthermore, heat map analysis deciphered the clustering of the gene expression profiles among the various datasets (Figure 6c). The results from the nCounter assays correlated with the transcriptomic analysis (Table 2, 3 and 4), hence substantiating the findings.

**Figure 6:**
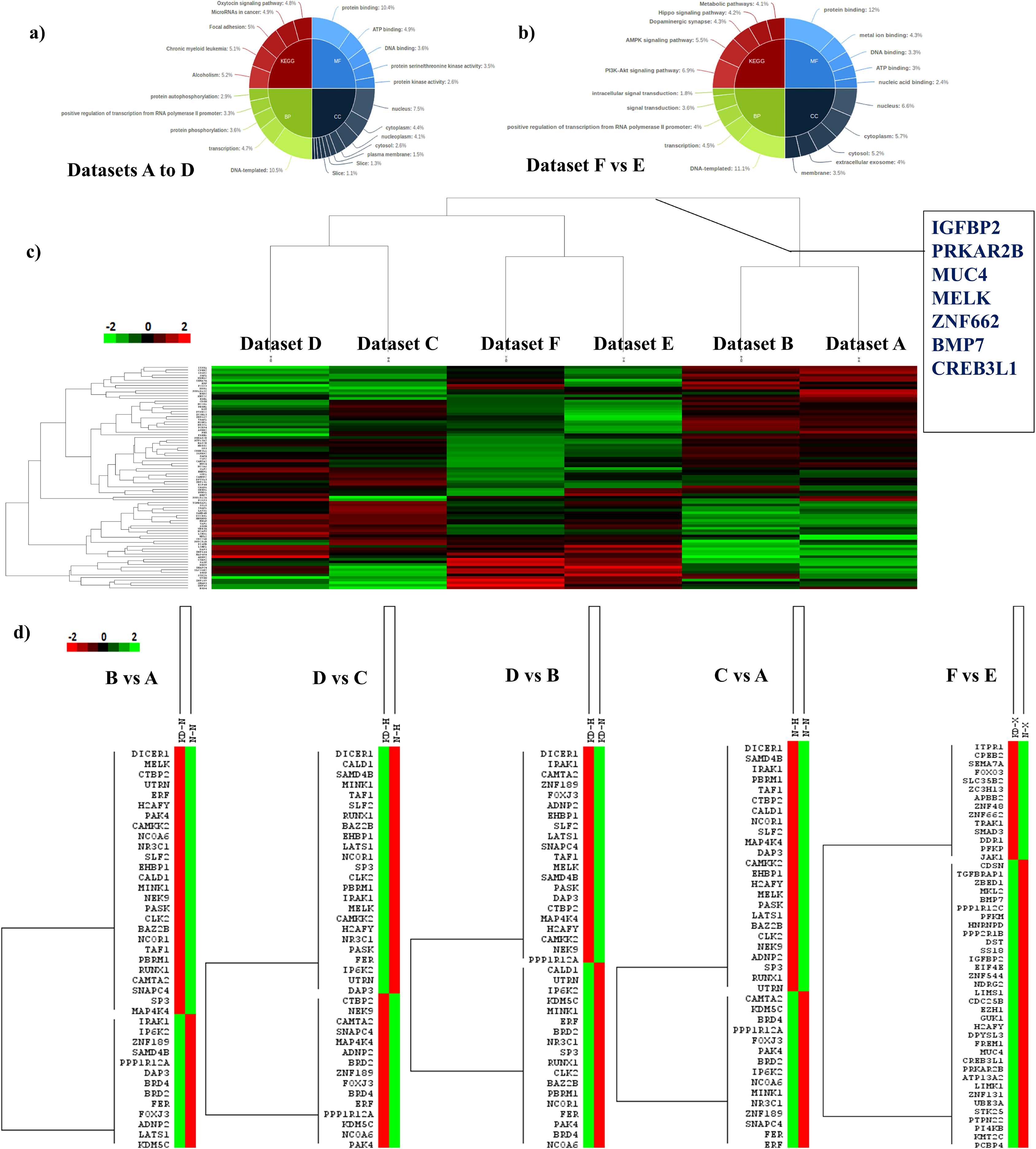
Pie chart representation of the top five gene ontologies and pathway summaries based on all the DEGs in the various analyzed datasets used for the nCounter gene expression assay for **a)** *in vitro* HNSCC cell line model (A-D datasets) and **b)** *in vivo* heterotopic HNSCC xenograft model (F vs E dataset). **c)** Representative nonhierarchical heatmap representation depicting the expression profile and variation in average log2 FC among the DEGs considering the complete CodeSet of 86 genes post-3’-UTR filtering and **d)** the individual CodeSet of 39 selected genes for the *in vitro* HNSCC cell line model (A-D datasets) and 48 selected genes for the *in vivo* heterotopic HNSCC xenograft dataset post-3’UTR-filtering. The various analyzed datasets used for expression profiling are labeled on the top: *in vitro* HNSCC cell line model (A-D datasets) where Lane 1-KDH is Dataset D (CAL27-MK2_KD_ cells in Hypoxia); Lane 2-NH is Dataset C (CAL27-MK2_WT_ cells in Hypoxia); Lane 5-KDN is Dataset B (CAL27-MK2_KD_ cells in Normoxia); Lane 6-NN is Dataset A (CAL27-MK2_WT_ cells in Normoxia), and *in vivo* heterotopic HNSCC xenograft dataset where Lane 3-KDX is CAL27-MK2_KD_ cells grafted (F dataset) and Lane 4-NX is CAL27-MK2_WT_ cells grafted (E dataset). Distinct clusters of upregulated and downregulated genes are visible in each combination. The color bar represents the expression values with green representing the upregulation and red representing the downregulation expression levels.

Individually, the expression profile and variation in FC among the DEGs in the nCounter analysis in the *in vitro* CodeSet of 39 genes (Table ST3 and ST7) and in the *in vivo* CodeSet of 48 genes (Table ST8) have been showcased *via* the heat-map representations in Figure 6d, respectively. The results depict the upregulated and downregulated DEGs in the nCounter assays and the results were in consonance with the transcriptomic profiling with a high percentage of genes showing a similar pattern of expression and even matching FC values as shown in Table 3 and 4. For the B vs A dataset, a total of 39 genes were analyzed, out of which 24 matched the transcriptomic analysis (61.6% matching score). The matched genes were then analyzed for FC and BRD2 was found to be the only upregulated gene (FC>2), while CLK2 was the only downregulated gene (FC<2). Similarly, a step-by-step comprehensive analysis of all the datasets was performed and 12 DEGs that were common in the transcriptomic and nCounter analysis were revealed (the results are summarized in Table 2). Notably, these genes are key players in important processes such as apoptosis, cell cycle progression, and transcription regulation, hence, potentiating their role as important MK2-regulated genes involved in HNSCC pathogenesis (Figure 7d). Therefore, these results strengthen our recent findings that MK2 is critically important in regulating HNSCC and functions by modulating the transcript stability of crucial genes driving pathogenesis. Furthermore, detailed statistical analysis accentuated that the expression of only 7 (BMP7, CREB3L1, IGFBP2, MELK, MUC4, PRKAR2B, and ZNF662), out of the 12 candidate genes were significantly different among the datasets (FC>2 or <-2, p<0.05) (Table 2, 3, and 4). MELK was the only gene belonging to the C vs A dataset (cell line comparison) while the other 6 genes were from the F vs E dataset (xenograft comparison). Moving forward, these 6 genes (BMP7, CREB3L1, IGFBP2, MUC4, PRKAR2B, and ZNF662) were analyzed *in vitro* by IHC and RT-qPCR analyses to ascertain the transcriptomic and nCounter findings in an experimental HNSCC xenograft model (Figure 7).

**Figure 7:**
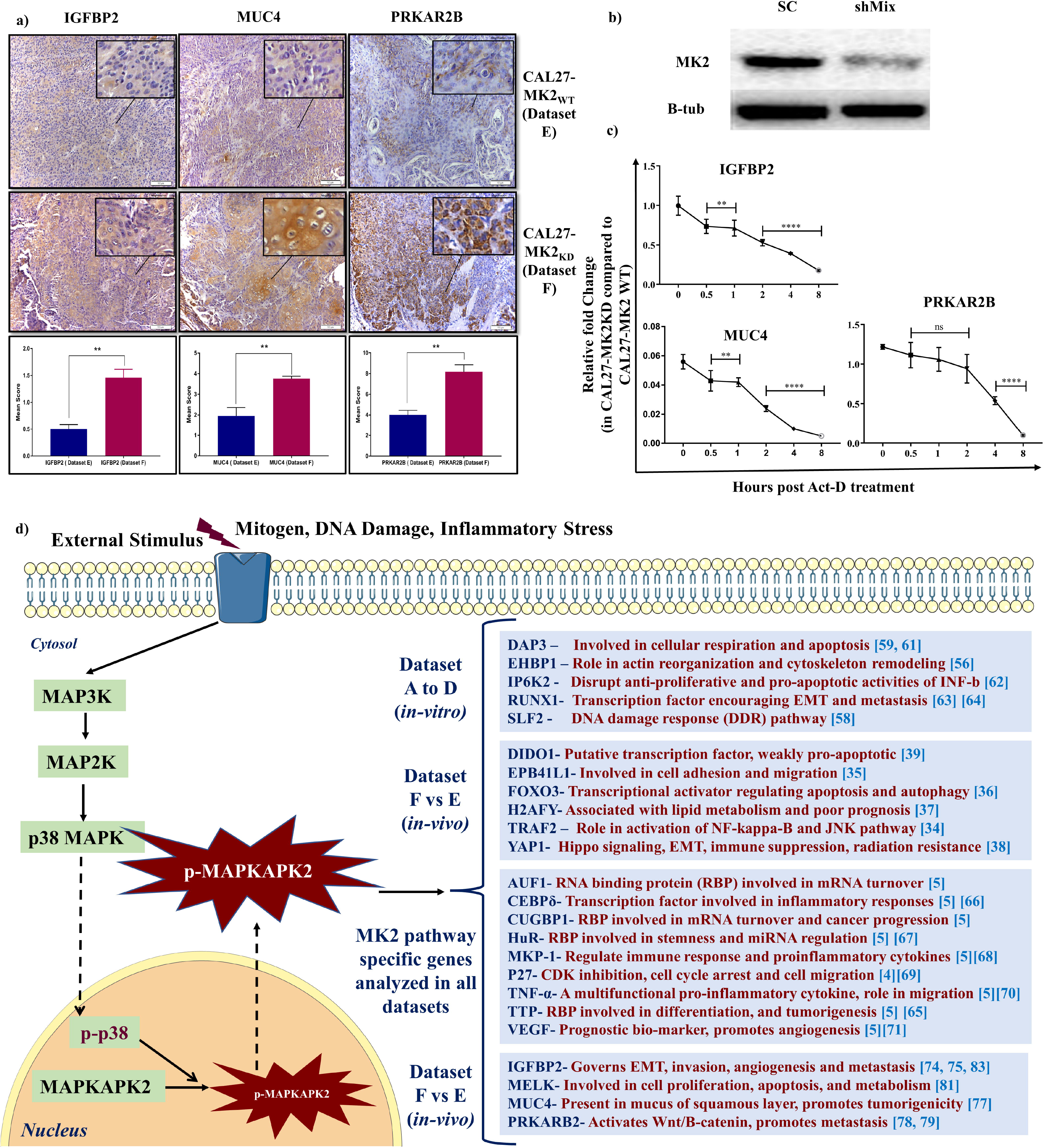
**a)** Representation of the *in-situ* protein expression levels of the three candidate MK2-regulated genes (IGFBP2, MUC4, PRKAR2B) in HNSCC xenografted mice tumor sections by immunohistochemical analysis. Different color bars represent the expression values in terms of mean score, the blue bar represents the expression level in CAL27-MK2_WT_ tissue sections (Dataset E), and the red bar signifies *In-situ* protein expression in CAL27-MK2_KD_ tissue sections (Dataset F). Parametric Welch t-test was used to evaluate the statistical significance using GraphPad Prism 7.0 software, ** denotes p<0.01; n=5 field views for IHC analysis. **b)** Western blot depicting MK2 expression in CAL27-MK2_KD_ cells generated post MK2 knockdown using shRNA for transcript stability experiment, here, SC-Scrambled control; shMix-A combination of an equal quantity of MK2 targeting shRNA complexes 1, 2, 3, and 4; the shMix group was used for transcript stability experiments, and SC was used as a control. **c)** Gene expression patterns of the 3 candidate genes in CAL27-MK2_KD_ cells at different time points post-Act-D treatment to assess the role of MK2 in regulating transcript stability. Data points represent the fold change in the shMix group at a particular time interval as compared to the control group. Linear regression analysis was performed using GraphPad Prism 7.0 software. **d)** Pictorial representation of the MAPK pathway analyzed in this study, the illustration depicts the activation of MK2 and the plausible mode of action in HNSCC pathogenesis. Figure elucidates the final MK2-regulated putative candidate genes in the pathway that were obtained in this study which could be further explored as possible targets for HNSCC management.

### 3.8 Immunohistochemical and RT-qPCR analyses indicated the putative role of MK2-regulated candidate genes in HNSCC pathogenesis

IHC and RT-qPCR were performed to probe the role of the 6 candidate genes in HNSCC pathogenesis and further accentuate the consonance among the transcriptomic and nCounter data analyses. Sections from the xenograft tumor tissues were analyzed using IHC to evaluate the protein expression pattern of the 6 candidate genes (BMP7, CREB3L1, IGFBP2, MUC4, PRKAR2B, and ZNF662) (Figure 7a). The results revealed that xenograft tumor sections with an unregulated expression of these genes have cellular pleomorphism, mitotic figures, and formation of nests of tumor cells, thus, clearly indicating the aggressiveness of HNSCC neoplasms. The results obtained from the IHC analysis largely strengthened the findings of the transcriptomic and nCounter analyses. Out of the 6 candidate genes, *in situ* protein expression levels of MK2-regulated candidate genes IGFBP2, MUC4, and PRKAR2B were found to be upregulated in the tumor xenografts created using CAL27-MK2_KD_ cells as compared to CAL27-MK2_WT_ cells and corroborated with the transcriptome and nCounter analyses (Figure 7a). There was no significant change in the protein expression levels of the 3 other analyzed genes BMP7, CREB3L1, and ZNF662 (data not shown).

Additionally, to examine the role of MK2 in governing the transcript stability of these genes, RT-qPCR analysis was performed in MK2-knockdown cells. CAL27 cells were treated with the shRNA complex to generate CAL27-MK2_KD_ cells as previously described [5] and shown in Figure 7b. Post-knockdown, CAL27-MK2_KD,_ and CAL27-MK2_WT_ (scrambled control transfected cells) were treated with Act-D (1 µM) to halt *de novo* transcription. RT-qPCR was carried out post-Act-D treatments at 6 different time points (0, 0.5, 1, 2, 4, and 8 hours) to determine the transcript expression levels and stability. Transcript levels were compared for each time point between the WT and KD group. The results revealed that the transcript expression of IGFBP2, MUC4, and PRKAR2B exhibited time-dependent transcriptional decay in CAL27-MK2_KD_ cells as compared to CAL27-MK2_WT_ cells (Figure 7c). The IGFBP2 transcript level was 0.99-fold in CAL27-MK2_KD_ cells which were equivalent to that in CAL27-MK2_WT_ cells (t=0), while the PRKAR2B transcript level was 1.2-fold higher in CAL27-MK2_KD_ cells as compared to CAL27-MK2_WT_ cells (t=0). The expression levels of these genes were stabilized for 1 hour for IGFBP2 and 2 hours for PRKAR2B post-Act-D treatment but gradually showed a significant reduction to less than half the levels (t=4 and t=8) in CAL27-MK2_KD_ cells compared to CAL27-MK2_WT_ cells (Figure 7c). This timeframe of stability may provide sufficient opportunity for the transcript to be expressed and upregulated at the protein level, which was observed in the IHC analysis of the tumor sections (Figure 7c). This suggested that the regulation could be at the transcript level and not at the protein level. However, the MUC4 transcript level was 16-fold lower in CAL27-MK2_KD_ cells as compared to CAL27-MK2_WT_ cells (t=0) but remained stable until 1 hour (t=1) post-Act-D treatment and decreased significantly (>50%) afterward (t=4) (Figure 7c). Taken together, the transcripts of all the 3 genes were degraded in MK2-knockdown cells after maintaining stable transcript levels for a few hours. This finding indicates a strong association of these genes with the expression profile of MK2 in the cells. Conclusively, the results suggested that the expression levels of IGFBP2, MUC4, and PRKAR2B are strongly affected by MK2 expression in HNSCC cells and tumors.

## 4. Discussion

To improve the understanding of convoluted biology and leverage the outcomes to optimize the management of HNSCC, there have been many efforts to characterize its pathology at the transcript level. Methodological breakthroughs in the recent past have revolutionized the area of transcriptome profiling by providing a link between molecular mechanisms and cellular phenotypes [23, 49]. In recent times, a comprehensive landscape of genomic and transcriptomic alterations in squamous tumors including HNSCC has emerged by way of the TCGA network [17, 18]. However, cellular models that can comprehensively characterize metastatic HNSCC are still lacking, hence, translationally relevant transcriptome profiling underlying the basis of HNSCC metastasis will prove to be a powerful tool for future preclinical research endeavors [50, 51]. help many established methods help in the detection of DEGs for both microarray-based approaches and RNA-seq [52, 53]. A typical transcriptome profiling result is generally a never-ending list comprising thousands of DEGs, hence, it has always been very difficult to interpret this data without additional filtering *via* functional annotations. A large variety of methods are available for the analysis of DEGs and for obtaining a critical understanding of the pathways, gene regulatory, and co-expression networks involved [54, 55]. In the present study, we undertook the challenge of thoroughly dissecting the huge complexity and large heterogeneity in HNSCC to discern novel biomarkers and potential therapeutic targets.

Keeping in mind the critical findings from previous studies, we performed transcriptome profiling of both the *in vitro* cell line as well as *in vivo* xenograft tumor samples that resulted in thousands of DEGs. These genes were segregated based on their clustering in various biological processes (Table ST2, ST4, and ST6). In line with the primary goal, the processes were filtered based on relevance in cancer leading us to 5 overlapping DEGs in the cell line datasets (A-D), viz. DAP3, EHBP, IP6K2, RUNX1, and SLF2, as listed in Table ST3. EHBP is encoded by the EHBP1 gene, and this protein has been shown to portray a role in actin reorganization and endocytic trafficking [56]. Polymorphism in this gene at the single nucleotide level has been reported to cause prostate cancer [57]. SLF2 is a DNA damage response pathway gene that functions by regulating genomic stability by post-replication repair of damaged DNA [58]. DAP3 has been shown to mediate interferon (IFN)-γ induced cell death in addition to its role in organelle biogenesis as well as maintenance and mitochondrial translation [59, 60]. DAP3 has been characterized by its pro-apoptotic function as a prognostic factor in gastric cancer and found to be associated with cancer progression [61]. The protein encoded by the IP6K2 gene has been shown to affect growth suppression and apoptotic action of IFN-β in the physiologic regulation of apoptosis in ovarian cancers with its deletion leading to HNSCC predisposition [62]. Lastly, the protein encoded by RUNX1 has been shown to be involved in the activation of EMT *via* the Wnt/β-catenin pathway and the promotion of metastasis in colon cancer [63]. Further, it has been reported that RUNX1 depletion in human HNSCC cells causes growth arrest [64]. Collectively, it is quite evident that all these MK2-regulated genes are playing a vital role in tumor pathogenesis, hence showing consistency with our previous finding of their involvement in HNSCC pathogenesis.

Similarly, a gene regulatory network was generated that depicted a detailed overview of the various inter-connections and the significantly enriched biological processes affected by the DEGs in the 14 cancer-specific biological processes in the transcriptome profiling of the *in vivo* heterotopic HNSCC xenograft dataset (F vs E comparison). TRAF2 has been reported to have a role in the activation of the NF-kappa-B and JNK pathways [34]. EPB41L1 has been shown to have a high prognostic significance and is involved in cell adhesion and migration [34 changed]. FOXO3 is a transcriptional activator known to regulate apoptosis and autophagy in various tumors [36]. H2AFY has been shown to be associated with lipid metabolism and poor prognosis in liver cancer [37]. YAP1 is an important candidate gene of the hippo signaling and has shown to be involved in EMT, immune suppression, and radiation resistance [38]. DIDO1 is a putative transcription factor and has been reported to have weak pro-apoptotic activities [39] (Figure 3b). In lieu of the above arguments and considering the role of the MK2-regulated candidate genes in the experimental analysis, their further exploration as candidates for the development of novel biomarkers and utilization as potential therapeutic targets in HNSCC management is warranted.

Additionally, gene regulatory networks in transcriptome profiling provided information on the various biological processes regulated by these candidate genes. This study supplied a wealth of information that can be further explored to study the pathogenesis of HNSCC in detail, especially in the background of MK2-knockdown and a varied tumor microenvironment (normoxia/hypoxia). Figure 4a is the representation of a gene regulatory network depicting the role of MK2 in the regulation of mRNA stability in the various datasets analyzed and clearly demonstrates that MK2 regulates transcript stability *via* RBP-mediated regulation with HuR (ELAVL1) and TTP (ZFP36) playing integral roles. Similarly, Figure 4b portrays the regulatory network that represents the MAPK signaling cluster of the selected MK2 pathway genes (AUF1, CEBP, CUGBP1δ, HuR, MK2, MKP-1, p27, p38, TNF-α, TTP, and VEGF) in the transcriptome profiling data of the *in vitro* HNSCC cell line dataset (B vs A, normoxic microenvironment) indicated TNF-α and VEGF downregulation. Interestingly, the transcriptomic profiling results are in complete consonance with our recently published findings and hence succeed in potentiating and validating the hypothesis that MK2-knockdown destabilized TNF-α and VEGF in normoxia *via* RBP-mediated interactions [5, 65, 66, 67, 68, 69, 70, 71, 72].

Transcriptome analysis techniques are commonly utilized in endeavors to decipher various molecular mechanisms of tumorigenesis and to fetch out novel prognostic and therapeutic markers [22, 23, 73]. In this study, we aimed to assess the MK2-regulated candidate genes playing prominent roles in HNSCC pathogenesis. Using the 3’-UTR-based filtering criterion detailed before, 34 genes in the *in vitro* A-D datasets and 48 genes in the xenograft dataset (listed in Table ST7 and ST8) were identified. Further validation using the nCounter gene expression assay system enabled the digital quantification and single-molecule imaging of multiple target RNA molecules using multicolor molecular barcodes (Figure S4 and S9). This system provides discrete and accurate counts of RNA transcripts at a high level of sensitivity and precision [32]. The gene expression assays are independent of any enzymatic reactions or amplification protocols and have no reliance on the degree of fluorescence intensity to determine target abundance. As a result of these characteristics, and the highly automated nature of barcoded sample processing, these assays result in highly accurate and reproducible outcomes. On average, approximately 52% matching score of transcriptome profiling data was obtained with nCounter gene expression assay-based validation which is considered a good percentage match considering the high-throughput nature of the analysis and the various datasets analyzed. Filtering of the DEGs in the matched data revealed a list of 12 genes (6 upregulated and 6 downregulated in the various analyzed datasets) that were common in the comprehensive nCounter system-based validation of transcriptomic profiling (Table 4). Intriguingly, these genes portray crucial roles in processes such as apoptosis (CLK2, MELK, MUC4), cell cycle regulation (CLK2, MELK), and transcription regulation (BRD2, H2AFY, SAMD4B, ZNF662).

Six candidate genes (BMP7, CREB3L1, IGFBP2, MUC4, PRKAR2B, and ZNF662) showed statistically significant up/downregulation in the xenograft dataset. Insulin-like Growth Factor Binding Protein 2 (IGFBP2) has been shown to be a growth promoter gene in several tumors and is considered a central hub of the oncogenic signaling network governing transcriptional regulation and promoting epithelial to mesenchymal transition, invasion, angiogenesis, and metastasis [74] Recently, IGFBP2 has been reported to be a crucial modulator of metastasis in oral cancer as well [75]. Mucin 4 (MUC4) serves as a major constituent of mucus secreted by epithelial cells and found overexpressed in a variety of cancers such as papillary thyroid carcinomas. It is known for promoting tumor growth, proliferation, and migration [76] [77]. Recent insights have been made into the transcriptional regulation of protein kinase cAMP-dependent type II regulatory subunit beta (PRKAR2B) by miRNAs and X-box binding protein 1 leading to a better understanding of PRKAR2B-driven prostate cancer progression [78]. PRKAR2B has been reported to be involved in the activation of Wnt/β□catenin along with triggering epithelial to mesenchymal transition leading to metastasis in tumors [79]. Consistent with our findings, these genes have been suggested to be prognostic indicators and therapeutic targets in various cancers including HNSCC [74, 75, 76, 77, 78, 79, 80, 81, 82].

The protein expression pattern of the 6 candidate genes was further analyzed in tumor sections from xenografted animals. IHC analysis revealed that IGFBP2, MUC4, and PRKAR2B were upregulated and prominently expressed in the cytoplasm and stroma of the tumors generated using CAL27-MK2_KD_ cells (Figure 7a). The expression levels of the other 3 genes were not significantly different among the samples. The IHC results were in consonance with the sequencing data, hence, confirming that these genes display differential expression patterns between CAL27-MK_WT_ and CAL27-MK2_KD_ sections. These genes are widely considered imperative to processes such as cell cycle progression, apoptosis, and transcriptional regulation. Various studies have reported their role as central hubs for cellular signaling during oncogenesis and modulating key cellular processes such as apoptosis, cell cycle progression, epithelial-mesenchymal transition, and metastasis [75, 76, 77, 78, 79]. Additionally, fewer studies have also reported the contrasting role of these genes in oncogenesis, which is influenced by various factors such as mutation and effects of other genes on the regulation of gene or protein expression of these three genes [83, 84].

Furthermore, RT-qPCR analysis was performed to quantify of transcript expression and behavior of these genes *in vitro* under MK_WT_ and MK2_KD_ conditions. CAL27-MK2_KD_ cells were treated with Act-D for different time points and mRNA transcript levels of IGFBP2, MUC4, and PRKAR2B were evaluated using transcript expression and stability analysis through RT-qPCR. It was observed that the stability of IGFBP2, PRKAR2B, and MUC4 transcripts decreased temporally in CAL27-MK2_KD_ cells as compared to the CAL27-MK2_WT_ cells. At t=0, the transcript level of IGFBP2 was at the basal level, MUC4 was downregulated while PRKAR2B was upregulated in CAL27-MK2_KD_ cells post-Act-D treatment. Furthermore, this decay increased significantly at t=1 for IGFBP2 and MUC4 and at t=2 for PRKAR2B (Figure 7c). The initial stability of transcripts could account for the upregulated protein expression observed in tumor sections of CAL27-MK2_KD_ xenografted mice as assessed by IHC (Figure 7a). This finding also indicates a strong dependence of IGFBP2, MUC4, and PRKAR2B2 on MK2-mediated regulation *via* RBP-activation/deactivation mechanism. Since these genes have previously been reported to be differentially expressed in tumor conditions, the present study substantiated the hypothesis suggesting the central role of MK2 in this molecular crosstalk. Additionally, these genes have been shown to be involved in a multitude of cellular processes such as the cell cycle, apoptosis, transcription, invasion, and metastasis. The contrasting gene and protein expression levels can be attributed to their regulation either at the transcriptional level or *via* post-translational modifications. This specific scientific question strongly warrants attention and future studies to delve deeper into the role of MK2-mediated activation and deactivation of RBPs that are involved in the transcript stability of these genes. Overall, the results obtained from IHC, and transcript stability analysis indicated the crucial role of MK2 in the modulation of the expression pattern of these genes in HNSCC tumors and cells. Finally, these findings clearly potentiate the importance of these MK2-regulated candidate genes in HNSCC pathogenesis.

Conclusively, the results suggested an observed dependence of these candidate genes on MK2 for their transcription in HNSCC cells and xenograft tumors. It is worth mentioning that all these genes are MK2-regulated and potentially play specific roles in HNSCC pathogenesis and progression. This suggests that they could potentially be used as putative candidates for further investigations regarding the design of molecular markers and therapeutics for HNSCC management. Hence, transcriptomic analysis followed by nCounter assay based primary validation, followed by IHC and transcript stability-based secondary validation has provided valuable findings that can aid in extending these observations in future HNSCC-targeted clinical and therapeutic exploratory research.

## 5. Conclusion

In conclusion, the present study substantiates the involvement of MK2 as a critically important factor in regulating HNSCC by modulating the transcript stability of downstream genes involved in pathogenesis. The probable mechanism of action is *via* RBP-mediated regulation and these results are in perfect consonance and augmentation with recent findings [5]. Comprehensively, few crucial MK2-regulated putative candidate genes were identified in this study, and their plausible involvement in HNSCC pathogenesis was elucidated, which could have further exploratory value as putative targets in HNSCC treatment and management (Figure 7). This study has made it possible to filter down from thousands of DEGs to a few potential candidate genes using comprehensive transcriptomic and *in vitro* validation approaches. To delve deeper into the clinical insights of these findings highlighting the role of MK2-mediated changes in HNSCC pathogenesis, the role of these 3 potential therapeutic targets warrants further detailed investigations for diagnostic and therapeutic interventions of HNSCC.

## 6. List of Abbreviations

3′-UTR: 3′-untranslated region
ActD: Actinomycin D
ARE(s): Adenylate-uridylate-rich element(s)
ATCC: American Type Culture Collection
CISBP: Catalog of Inferred Sequence Binding Preferences
Ct: Cycle Threshold
DEG(s): Differentially expressed gene(s)
EHBP1: EH domain binding protein 1
SLF2: SMC5-SMC6 complex localization factor 2
DAP3: Death associated protein 3
FC: Fold change
FDR: False discovery rate
FPKM: Fragments per kilobase of transcript per million mapped
GFP: Green fluorescent protein
GO: Gene Ontology
HKG: House keeping genes
HNSCCs: Head and neck squamous cell carcinoma(s)
HQ: High quality
IAEC: Institutional animal ethics committee
IFN: Interferon
IGFBP2: Insulin-like growth factor-binding protein 2
IP6K2: Inositol hexakisphosphate kinase 2
IHC: Immunohistochemistry
KD: Knockdown
KEGG: Kyoto encyclopedia of genes and genomics
MAPK: Mitogen-Activated Protein Kinase
MAPKAPK2 or MK2: Mitogen-activated protein kinase-activated protein kinase 2
MK2_KD_: MK2-knockdown
MK2_WT_: MK2 wild-type
MELK: Maternal embryonic leucine zipper kinase
MUC4: Mucin 4
MKP-1: Mitogen-activated protein kinase phosphatase-1
NGS: Next generation sequencing
NOD/SCID: Non-obese diabetic/severe combined immunodeficient
p27: Cyclin-dependent kinase inhibitor 1B
PRKAR2B: Protein kinase CAMP-dependent type II regulatory subunit beta
QC: Quality control
RIN: RNA integrity number
RT-qPCR: Real-time quantitative polymerase chain reaction
RBP(s): RNA-binding protein(s)
RNA-seq: Ribose Nucleic Acid -sequencing
RUNX1: Runt-related transcription factor 1 shRNA Short hairpin RNA
TCGA: The cancer genome atlas
TNF-α: Tumor necrosis factor-alpha
TTP: Tristetraprolin
VEGF: Vascular endothelial growth factor
WB: Western blotting
WT: Wild type
ZNF662: Zinc finger protein 662

## 7. Declarations

### 7.1 Availability of Data and materials

The generated datasets (raw reads) from NovaSeq 6000 have been deposited and are available in the National Centre for Biotechnology Information-Sequence Read Archive (NCBI-SRA) repository (https://www.ncbi.nlm.nih.gov/sra). The SRA BioProject accession numbers for the submitted bio projects are PRJNA646850 and PRJNA646851. High-resolution images will be available on request if needed.

### 7.4 Authors’ contributions

YSP and SS conceptualized, designed the work, and framed the manuscript. SS and PA performed bench work, experiments, and data analysis. MKS helped with RNA-seq and data analysis, VP performed IHC imaging and analysis, and NVT helped in xenograft model generation. SS, PA, VP, and YSP wrote and edited the manuscript. All the authors read and approved the final version of the manuscript.

## 7.5 Acknowledgments

The authors would like to thanks to the Director, CSIR-IHBT, Palampur for his consistent encouraging support. SS is immensely thankful to CSIR and PA to DST (DST/INSPIRE Fellowship/2017/IF170140) for providing Ph.D. fellowship and Academy of Scientific and Innovative Research (AcSIR), Ghaziabad, India for Ph.D. registration. The authors would like to thank Bionivid Technology Private Limited, Bengaluru, India for their inputs in transcriptome data analysis. CSIR-IHBT communication number of this manuscript is 4682.

## 7.6 Additional Files

There are 2 Additional files for this manuscript: Additional File 1 (9 supplementary figures, labeled as Figure S1 – Figure S9) and Additional File 2 (10 supplementary tables, labeled as Figure ST1 – Figure ST10).

