## Supplementary material for "RNA-sequencing manifests the intrinsic role of MAPKAPK2 in facilitating molecular crosstalk during HNSCC pathogenesis": Tables

**Table 1: Sample description and comparison details.**

| **S. No.** | **Analysis Code** | **Sample Type** | **Sample Description** |
| --- | --- | --- | --- |
| 1. | A | Cultured Cell Line  (*in-vitro*) | Normal CAL27 cells (Normoxia) |
| 2. | B |  | MK2-knockdown CAL27 cells (Normoxia) |
| 3. | C |  | Normal CAL27 cells (Hypoxia) |
| 4. | D |  | MK2-knockdown CAL27 cells (Hypoxia) |
| 5. | E | Dissected Xenografts  (*in vivo*) | Normal CAL27 cells grafted |
| 6. | F |  | MK2-knockdown CAL27 cells grafted |
| **Comparison/**  **Dataset Code** | | **Comparison/Dataset Detail** | |
| **B vs A** **or**  CAL27-MK2_KD_ (N) vs CAL27-MK2_WT_ (N) | | MK2-knockdown CAL27 cells (Normoxia) vs  Normal CAL27 cells (Normoxia) | |
| **D vs C** **or**  CAL27-MK2_KD_ (H) vs CAL27-MK2_WT_ (H) | | MK2-knockdown CAL27 cells (Hypoxia) vs  Normal CAL27 cells (Hypoxia) | |
| **D vs B** **or**  CAL27-MK2_KD_ (H) vs CAL27-MK2_KD_ (N) | | MK2-knockdown CAL27 cells (Hypoxia) vs  MK2-knockdown CAL27 cells (Normoxia) | |
| **C vs A** **or**  CAL27-MK2_WT_ (H) vs CAL27-MK2_WT_ (N) | | Normal CAL27 cells (Hypoxia) vs  Normal CAL27 cells (Normoxia) | |
| **F vs E** **or**  CAL27-MK2_KD_ (X) vs CAL27-MK2_WT_ (X) | | MK2-knockdown CAL27 cells grafted vs  Normal CAL27 cells grafted | |

Tabular representation of the codes and details of all the samples used for transcriptome analysis and details of the various experimental datasets (comparisons) used in the study.

**Table 2: Summary of the comparison**

| **Combination** | **No. of genes analyzed** | **No. and % of genes matching with transcriptome analysis** | **Upregulated genes** | **Downregulated genes** |
| --- | --- | --- | --- | --- |
| **B vs A** | 39 | **24** (61.6 %) | **1** (BRD2) | **1** (CLK2) |
| **D vs C** | 39 | **17** (43.6%) | - | - |
| **D vs B** | 39 | **19** (48.7%) | - | **1** (SAMD4B) |
| **C vs A** | 39 | **20** (51.3%) | - | **2** (H2AFY, **MELK^*^**) |
| **F vs E** | 48 | **26** (54.2%) | **5** (**BMP7^*^, CREB3L1^*^, IGFBP2^*^, MUC4^*^, PRKAR2B^*^)** | **2** (CDSN, **ZNF662^*^**) |

Tabular representation of the summary of the comparison between the data obtained from transcriptome profiling and nCounter gene expression assay analysis for all the experimental datasets. Here, only 7 (shown by asterisk ‘*’ and highlighted in bold), out of the 12 candidate genes were significantly different among the datasets (FC>2 or <-2, p<0.05) and taken further for validations.

**Table 3: List of 39 genes from *in-vitro* model**

| **Gene Symbol** | **Entrez**  **GeneID** | **Gene name** | **C vs A** | **C vs A Nanostring** |
| --- | --- | --- | --- | --- |
| ADNP2 | 22850 | ADNP homeobox 2 | 0.26 | -1.10 |
| BAZ2B | 29994 | Bromodomain adjacent to zinc finger domain 2B | -0.28 | -1.12 |
| BRD2 | 6046 | Bromodomain containing 2 | - | 1.71 |
| BRD4 | 23476 | Bromodomain containing 4 | 0.39 | 1.58 |
| CALD1 | 800 | Caldesmon 1 | -0.24 | -1.31 |
| CAMKK2 | 10645 | Calcium/calmodulin dependent protein kinase kinase 2 | 0.78 | -1.74 |
| CAMTA2 | 23125 | Calmodulin binding transcription activator 2 | 0.24 | 1.07 |
| CLK2 | 1196 | CDC like kinase 2 | -2.71 | -1.19 |
| CTBP2 | 1488 | C-terminal binding protein 2 | -0.14 | -1.15 |
| DAP3 | 7818 | Death associated protein 3 | -1.49 | -1.92 |
| DICER1 | 23405 | Dicer 1, ribonuclease III | 2.57 | -2.21 |
| EHBP1 | 23301 | EH domain binding protein 1 | -3.28 | -1.34 |
| ERF | 2077 | ETS2 repressor factor | -0.04 | 1.21 |
| FER | 2241 | FER tyrosine kinase | -0.75 | 1.09 |
| FOXJ3 | 22887 | Forkhead box J3 | 1.18 | 1.20 |
| H2AFY | 9555 | H2A histone family member Y | -4.65 | -1.95 |
| IP6K2 | 51447 | Inositol hexakisphosphate kinase 2 | 0.55 | 1.71 |
| IRAK1 | 3654 | Interleukin 1 receptor associated kinase 1 | -8.54 | -1.93 |
| KDM5C | 8242 | Lysine-specific demethylase 5C | 0.25 | 1.40 |
| LATS1 | 9113 | Large tumor suppressor kinase 1 | -0.66 | -1.27 |
| MAP4K4 | 9448 | Mitogen-activated protein kinase kinase kinase kinase 4 | 4.16 | -1.36 |
| MELK | 9833 | Maternal embryonic leucine zipper kinase | -3.68 | -4.59 |
| MINK1 | 50488 | Misshapen like kinase 1 | -0.24 | 1.37 |
| NCOA6 | 23054 | Nuclear receptor coactivator 6 | - | 1.11 |
| NCOR1 | 9611 | Nuclear receptor corepressor 1 | -0.30 | -1.05 |
| NEK9 | 91754 | NIMA related kinase 9 | 5.34 | -1.10 |
| NR3C1 | 2908 | Nuclear receptor subfamily 3 group C member 1 | -0.15 | 1.28 |
| PAK4 | 10298 | P21 (RAC1) activated kinase 4 | - | 1.03 |
| PASK | 23178 | PAS domain containing serine/threonine kinase | - | -1.99 |
| PBRM1 | 55193 | Protein polybromo-1 | - | -1.37 |
| PPP1R12A | 4659 | Protein phosphatase 1 regulatory subunit 12A | 1.00 | 1.10 |
| RUNX1 | 861 | Runt related transcription factor 1 | 0.47 | -1.03 |
| SAMD4B | 55095 | Sterile alpha motif domain containing 4B | -2.54 | -1.52 |
| SLF2 | 55719 | SMC5-SMC6 complex localization factor 2 | -0.81 | -1.52 |
| SNAPC4 | 6621 | Small nuclear RNA activating complex polypeptide 4 | 7.41 | 1.09 |
| SP3 | 6670 | Sp3 transcription factor | 0.14 | -1.02 |
| TAF1 | 6872 | TATA-box binding protein associated factor 1 | -0.76 | -1.82 |
| UTRN | 7402 | Utrophin | -9.26 | -1.00 |
| ZNF189 | 7743 | Zinc finger protein 189 | -1.01 | 1.30 |

Tabular representation of the list of 39 genes in *in vitro* HNSCC cell line model (C vs A dataset) showing the match of the average log2 fold change values in the last two columns in both transcriptome profiling and the nCounter gene expression assay (NanoString gene expression assay). The average log2 fold change values are colored according to gene expression changes (red indicates upregulation while green indicates downregulation). The gene highlighted in blue is the matched candidate DEG that shows a statistically significant change in expression.

**Table 4: List of 48 genes from *in-vivo* xenograft model**

| **Gene Symbol** | **Entrez**  **GeneID** | **Gene name** | **F vs E** | **F vs E Nanostring** |
| --- | --- | --- | --- | --- |
| APBB2 | 323 | Amyloid beta precursor protein binding family B member 2 | -3.74 | -1.16 |
| ATP13A2 | 23400 | ATPase 13A2 | 3.75 | 1.45 |
| BMP7 | 655 | Bone morphogenetic protein 7 | 5.95 | 47.09 |
| CDC25B | 994 | Cell division cycle 25B | 0.73 | 1.18 |
| CDSN | 1041 | Corneodesmosin | -2.35 | 1.72 |
| CPEB2 | 132864 | Cytoplasmic polyadenylation element binding protein 2 | -3.07 | -1.14 |
| CREB3L1 | 90993 | Camp responsive element binding protein 3 like 1 | 3.46 | 15.34 |
| DDR1 | 780 | Discoidin domain receptor tyrosine kinase 1 | 6.31 | -1.18 |
| DPYSL3 | 1809 | Dihydropyrimidinase like 3 | - | 16.6 |
| DST | 667 | Dystonin | -6.97 | 1.09 |
| EIF4E | 1977 | Eukaryotic translation initiation factor 4E | 3.08 | 1.1 |
| EZH1 | 2145 | Enhancer of zeste 1 polycomb repressive complex 2 subunit | - | 1.37 |
| FOXO3 | 2309 | Forkhead box O3 | -7.23 | -1.5 |
| FREM1 | 158326 | FRAS1 related extracellular matrix 1 | -5.95 | 1.51 |
| GUK1 | 2987 | Guanylate kinase 1 | 0.11 | 1.2 |
| H2AFY | 9555 | H2A histone family member Y | 3.77 | 1.34 |
| HNRNPD | 3184 | Heterogeneous nuclear ribonucleoprotein D | -1.10 | 1.29 |
| IGFBP2 | 3485 | Insulin like growth factor binding protein 2 | 3.86 | 12.21 |
| ITPR1 | 3708 | Inositol 1,4,5-trisphosphate receptor type 1 | 7.67 | -1.11 |
| JAK1 | 3716 | Janus kinase 1 | -0.08 | -1 |
| KMT2C | 58508 | Lysine methyltransferase 2C | 8.64 | 1.1 |
| LIMK1 | 3984 | LIM domain kinase 1 | -4.37 | 1.26 |
| LIMS1 | 3987 | LIM zinc finger domain containing 1 | 6.28 | 1.27 |
| MKL2 | 57496 | MKL1/myocardin like 2 | 2.40 | 1.14 |
| MUC4 | 4585 | Mucin 4, cell surface associated | 2.82 | 7.74 |
| NDRG2 | 57447 | NMYC downstream-regulated gene 2 | - | 2.72 |
| PCBP4 | 57060 | Poly(RC) Binding Protein 4 | - | 1.07 |
| PFKM | 5213 | Phosphofructokinase, Muscle | 1.00 | 2.62 |
| PFKP | 5214 | Phosphofructokinase, platelet | -5.36 | -1.09 |
| PI4KB | 5298 | Phosphatidylinositol 4-kinase beta | 4.87 | 1.05 |
| PPP1R12C | 54776 | Protein phosphatase 1 regulatory subunit 12C | -4.14 | 1.12 |
| PPP2R1B | 5519 | Protein phosphatase 2 scaffold subunit Abeta | -5.42 | 1.03 |
| PRKAR2B | 5577 | Protein kinase camp-dependent type II regulatory subunit beta | 2.06 | 4.28 |
| PTPN22 | 26191 | Protein tyrosine phosphatase, non-receptor type 22 | 0.54 | 1.09 |
| SEMA7A | 8482 | Semaphorin 7A (John Milton Hagen blood group) | - | -1.74 |
| SLC35B2 | 347734 | Solute carrier family 35 member B2 | 1.57 | -1.01 |
| SMAD3 | 4088 | SMAD family member 3 | 0.00 | -1.29 |
| SS18 | 6760 | SS18, nbaf chromatin remodeling complex subunit | -4.34 | 1.25 |
| STK25 | 10494 | Serine/threonine kinase 25 | - | 1.15 |
| TGFBRAP1 | 9392 | Transforming growth factor beta receptor associated protein 1 | 3.62 | 1.22 |
| TRAK1 | 22906 | Trafficking kinesin protein 1 | - | -1.26 |
| UBE3A | 7337 | Ubiquitin-protein ligase E3A | -0.06 | 1.14 |
| ZBED1 | 9189 | Zinc finger BED-type containing 1 | -3.63 | 1.68 |
| ZC3H13 | 23091 | Zinc finger CCCH-type containing 13 | 0.27 | -1.03 |
| ZNF131 | 7690 | Zinc finger protein 131 | - | 1.26 |
| ZNF48 | 197407 | Zinc finger protein 48 | -0.98 | -3.04 |
| ZNF544 | 27300 | Zinc finger protein 544 | - | 1.16 |
| ZNF662 | 389114 | Zinc finger protein 662 | -3.48 | -3.44 |

Tabular representation of the list of 48 genes in the *in vivo* heterotopic HNSCC xenograft experimental dataset (F vs E dataset) showing the match of the average log2 fold change values in the last two columns in both transcriptome profiling and the nCounter gene expression assay (NanoString gene expression assay). The average log2 fold change values are colored according to gene expression changes (red indicates upregulation while green indicates downregulation). The 6 highlighted genes in blue are the matched candidate DEGs that show a statistically significant change in expression.
