## Supplemental Figures for "RNA-sequencing manifests the intrinsic role of MAPKAPK2 in facilitating molecular crosstalk during HNSCC pathogenesis"

#### Additional File 1

**Authors' information**

[1]Pharmacology and Toxicology Laboratory, CSIR-Institute of Himalayan Bioresource Technology (CSIR-IHBT), Palampur-176061, India

[2]Biotechnology Division, CSIR-Institute of Himalayan Bioresource Technology (CSIR-IHBT), Palampur-176061, India

[3]Academy of Scientific and Innovative Research (AcSIR), Ghaziabad-201002, India.

**Figures:**

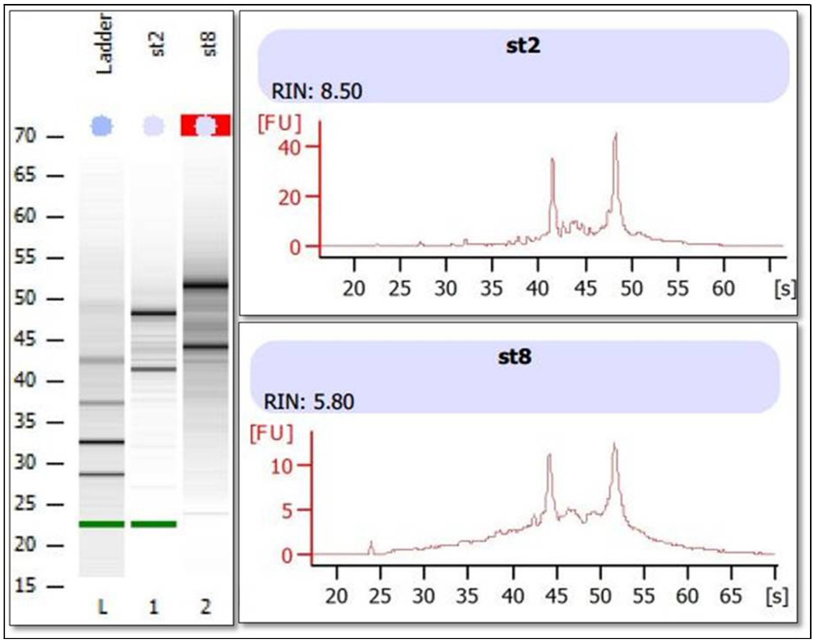


**Figure S1:** RNA quality assessment using Bioanalyzer showing comparative intensities of RNA bands from two representative samples used in the study (left); and electropherograms of both the samples (right) depicting intensity of intact RNA along with their RIN values (8.5 and 5.8) suggesting good quality RNA.

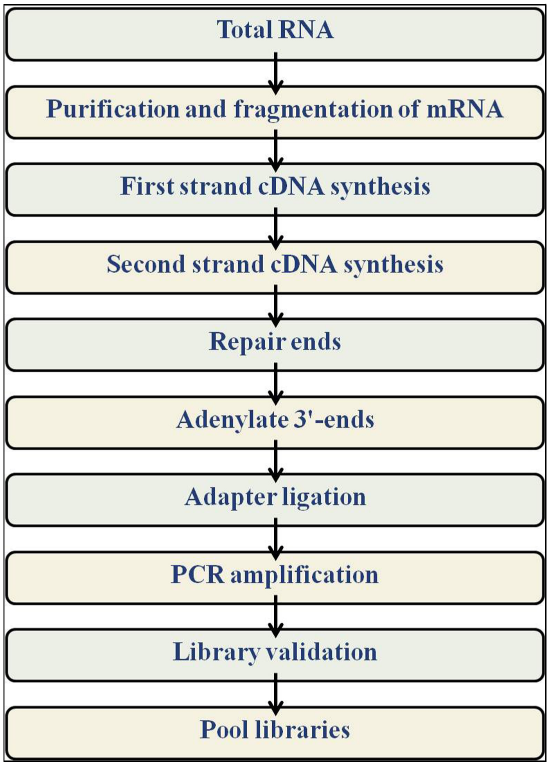


**Figure S2:** Detailed workflow of the process of library preparation using Illumina TruSeq RNA sample prep kit v2.

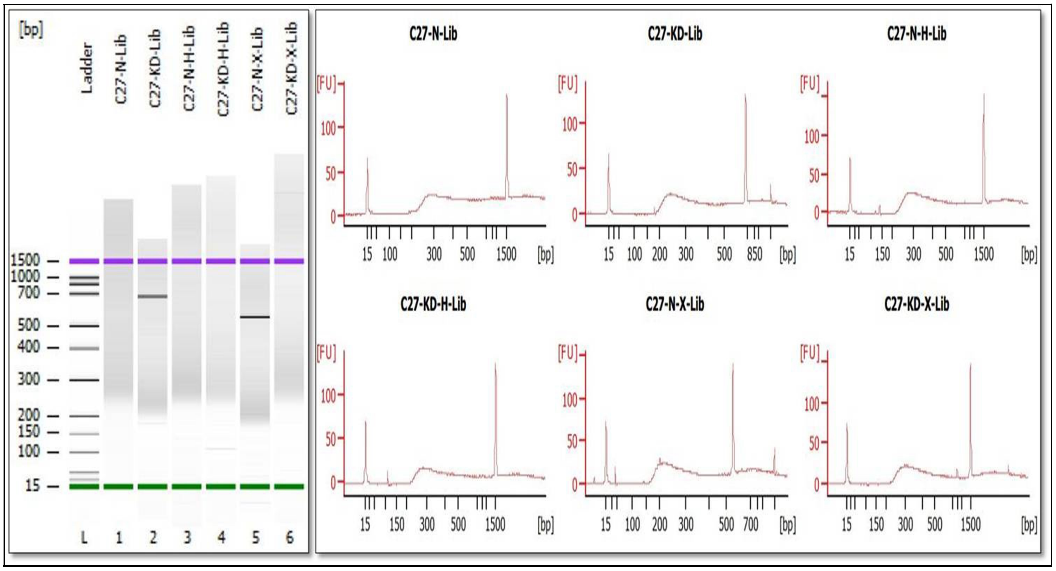


**Figure S3:** Bioanalyzer based quality check of all the six cDNA libraries used for sequencing showing comparative intensities of cDNA bands (left) and respective electropherograms (right).

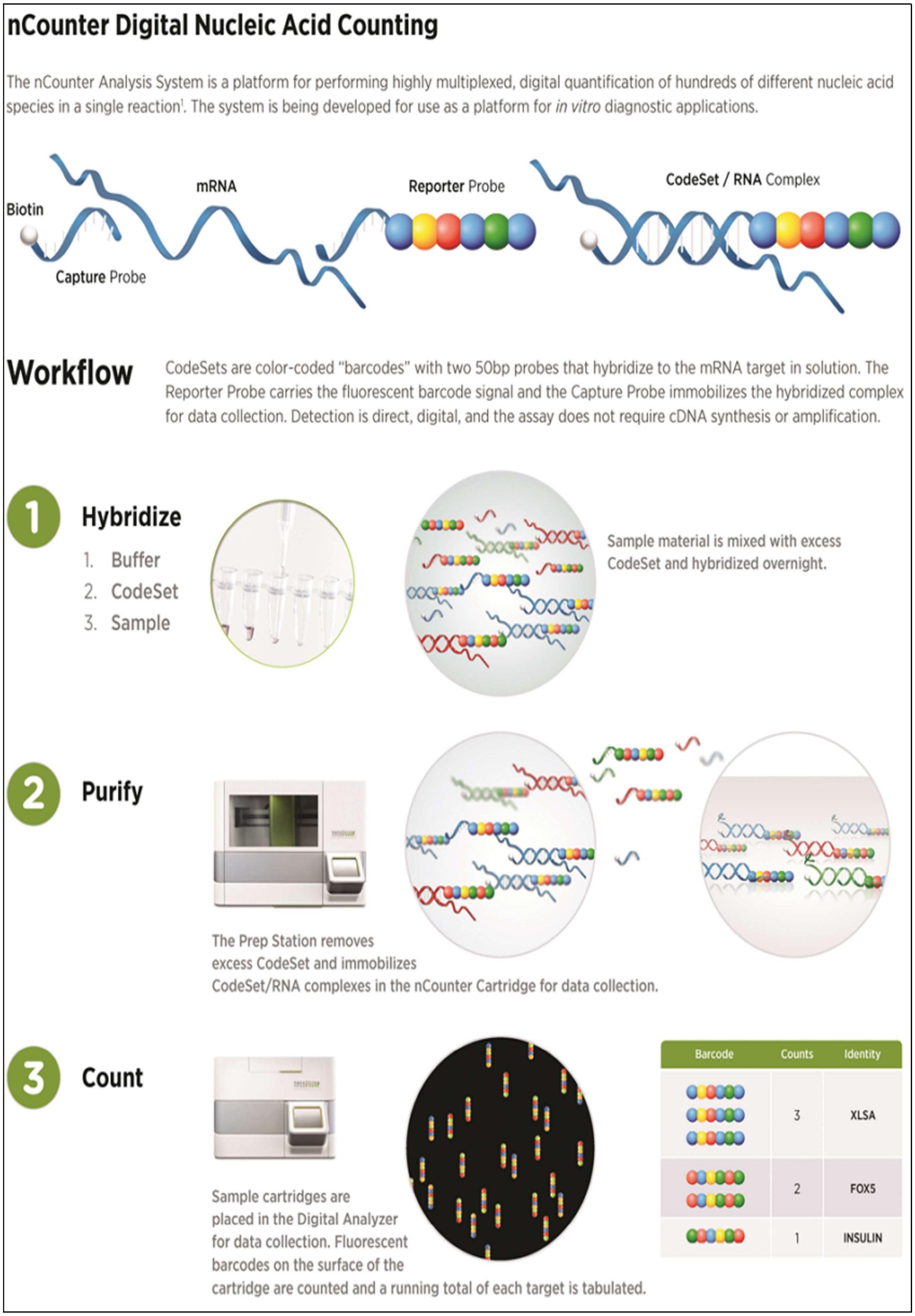


**Figure S4:** A comprehensive schematic representation of the workflow of nCounter digital nucleic acid counting which forms the basis for the nCounter gene expression assay (Source: NanoString Technologies, USA).

### Experiment QC Report

**Sequencing Run Details:**

- No. of Samples: 06
- Sample Type: human cell lines
- Name of the Instrument: Illumina NovaSeq 6000
- Read length: 100 bp (2X100bp)
- Run Type: Paired End
- Average library insert size: 210 bp

**Detailed table:**

| S.No. | Samples/ Conditions | No. of raw reads | No. of filtered reads | % | Total bases generated |
|---|---|---|---|---|---|
| 1 | A | 66,214,847 | 50,376,579 | 76.08 | 5,004,115,297 |
| 2 | B | 55,837,013 | 39,909,254 | 71.48 | 3,963,335,777 |
| 3 | C | 52,710,517 | 39,369,190 | 74.69 | 3,910,033,984 |
| 4 | D | 63,572,238 | 47,365,561 | 74.51 | 4,704,086,013 |
| 5 | E | 52,009,268 | 39,849,826 | 76.62 | 3,957,452,968 |
| 6 | F | 59,022,920 | 42,018,319 | 71.19 | 4,174,074,283 |
| | Total | 349,366,803 | 258,888,729 | 74.09 (Ave) | 25,713,098,322 |

- **Total no. raw reads: 349 Millions**
- **Total filtered reads: 258 Millions (filtered with NGS-QC Tool kit)**
- **Total generated sequencing data: 25.7 Gbases**

**Figure S5:** Tabular representation of the experimental quality check report for the sequencing run showing the details of the raw and filtered reads generated for all the experimental samples.

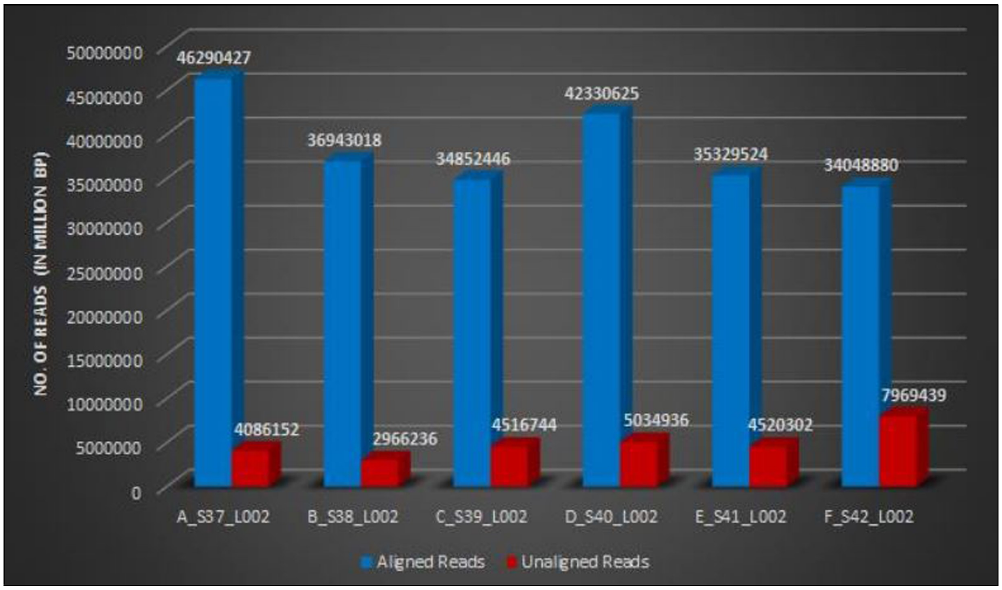


**Figure S6:** Bar diagram representation of the number of high-quality reads that could be mapped to the human reference genome in all the experimental datasets for the transcriptome profiling (blue bar indicates alignment while red bar shows unaligned reads).

a)

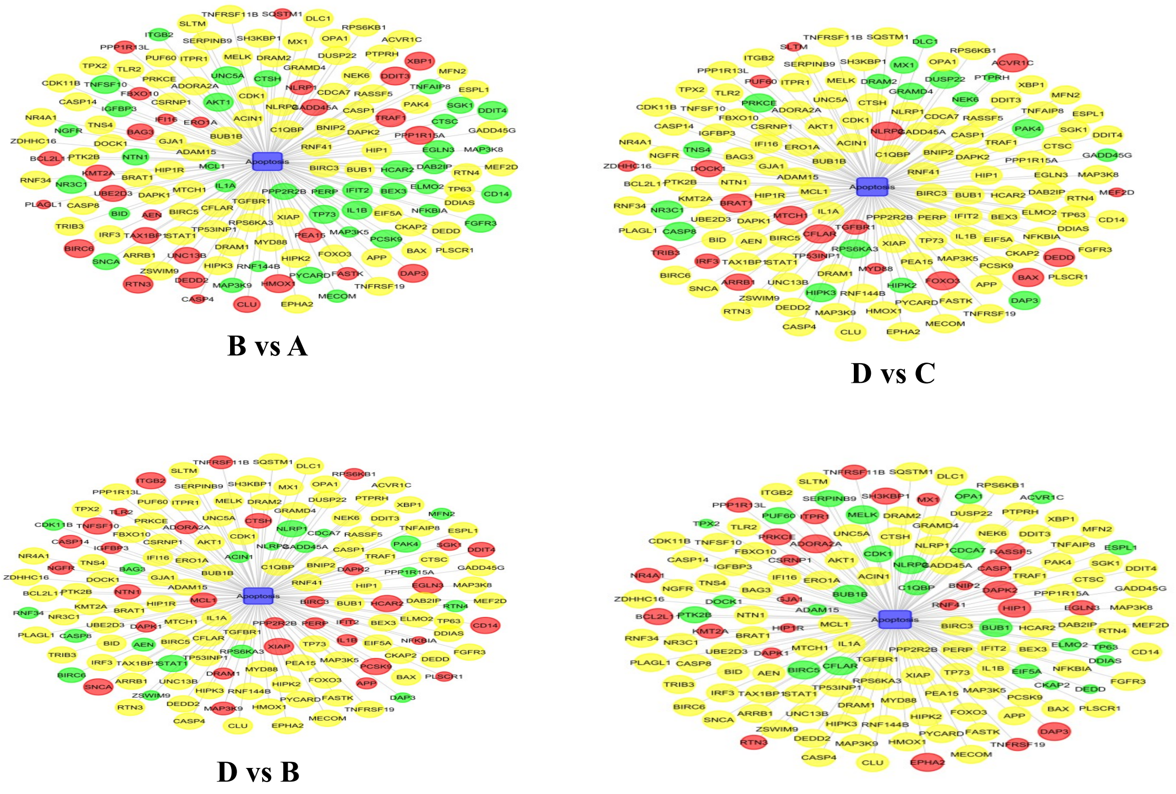

B vs A

D vs C

D vs B

C vs A

b)

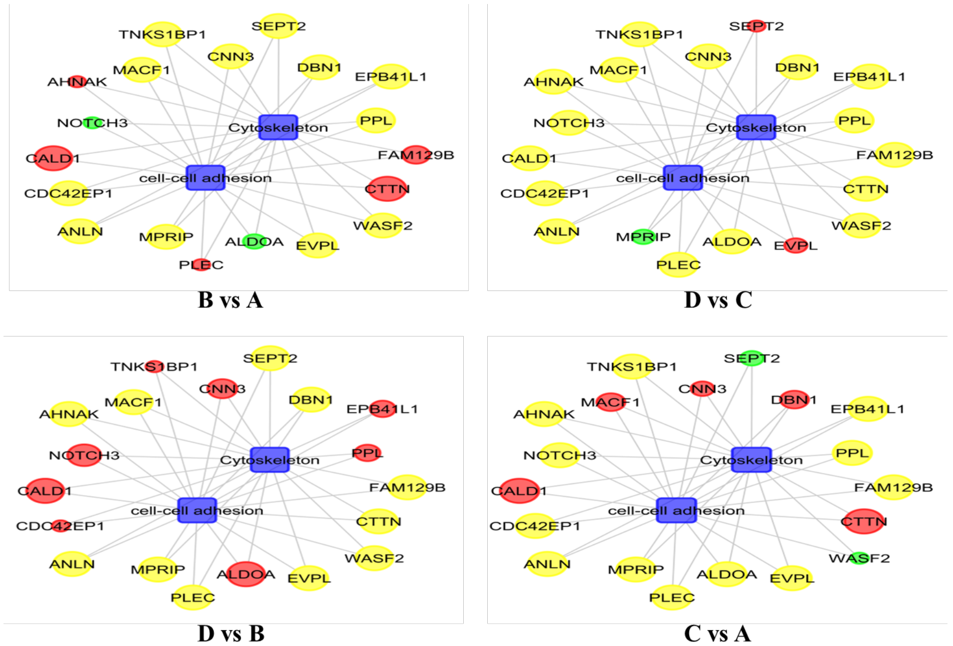


B vs A

D vs C

D vs B

C vs A

c)

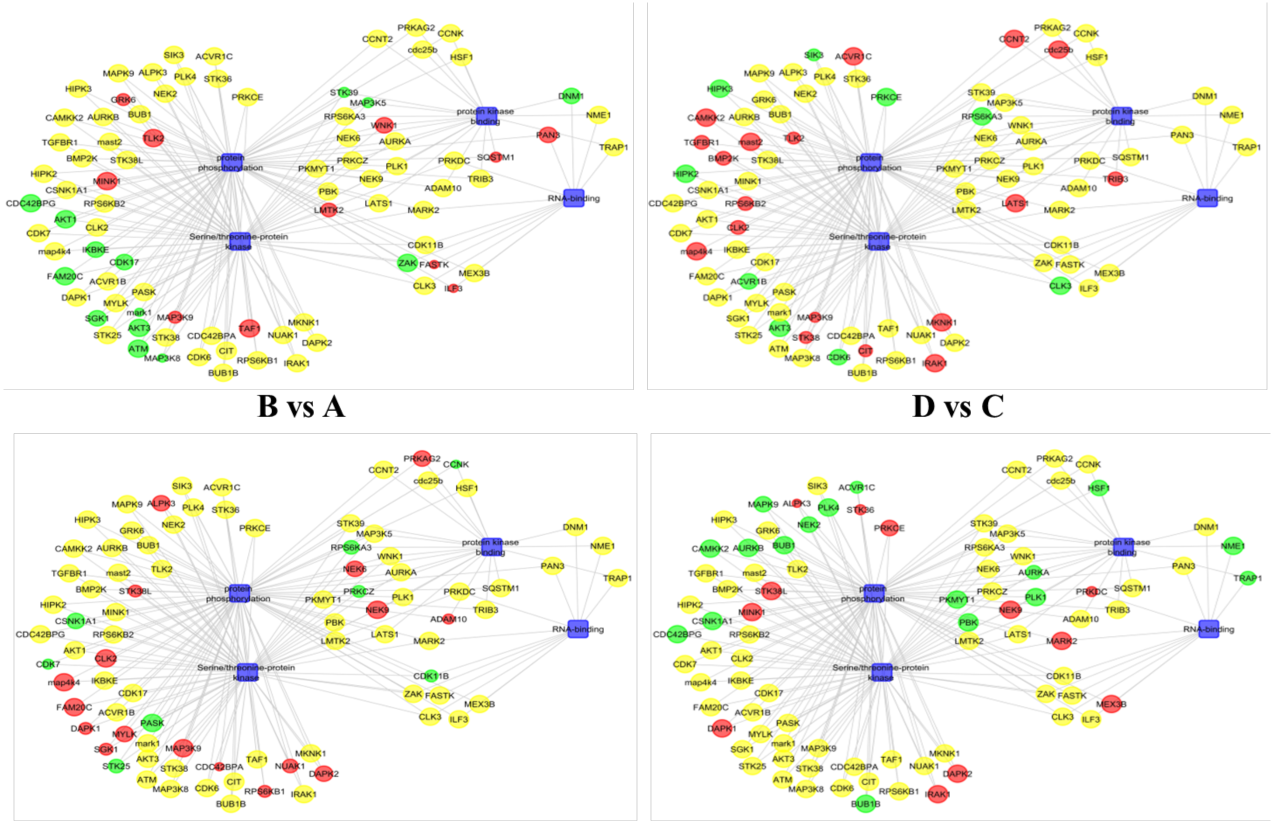


**B vs A**

**D vs C**

**D vs B**

**C vs A**

d)

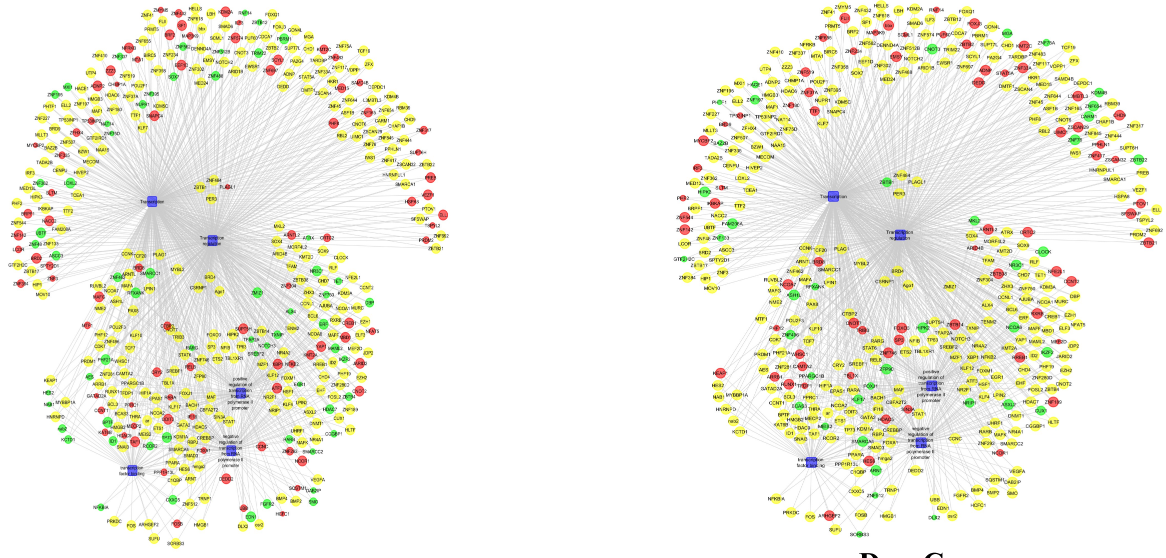

**B vs A**

**D vs C**

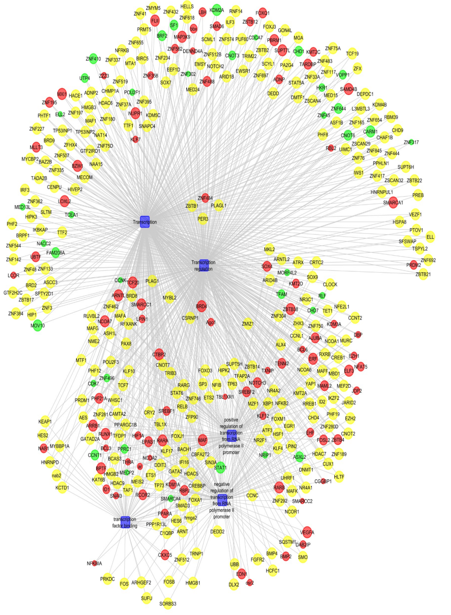

**D vs B**

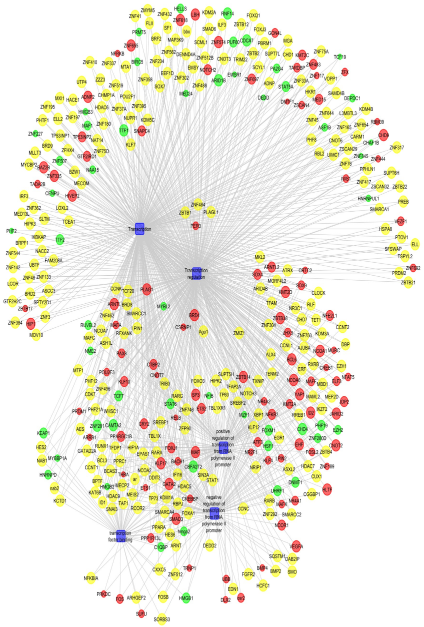

**C vs A**

**e)**

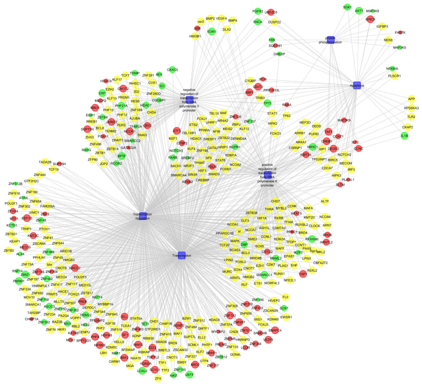

**B vs C**

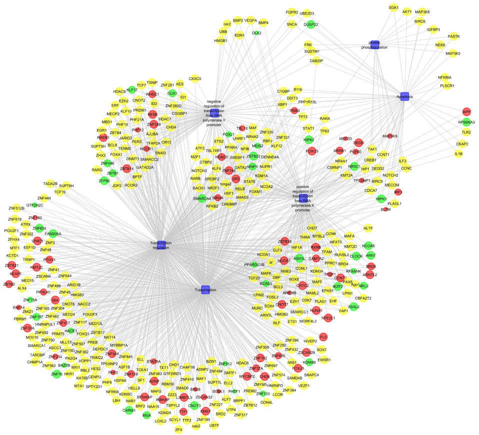

**D vs C**

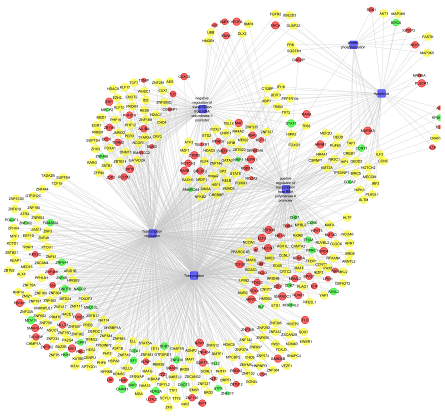

**D vs B**

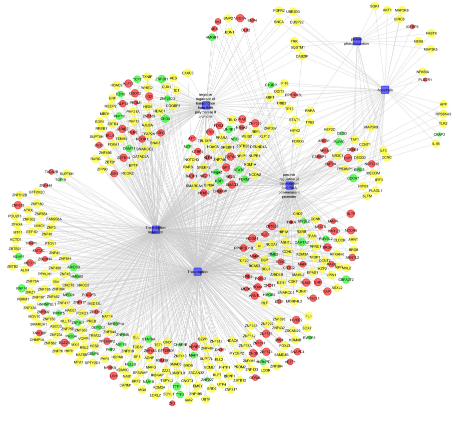

**C vs A**

**Figure S7:** Clustering of the 16 common elements in the transcriptome profiling of the *in vitro* HNSCC cell line model (A-D comparisons) into five major biological pathways.

**a)**

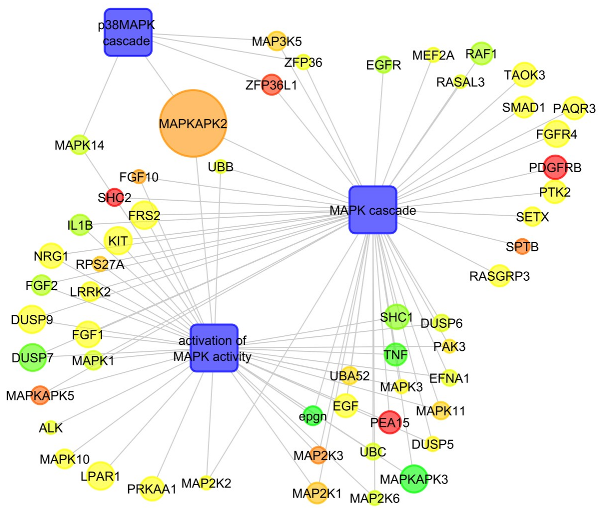


**b)**

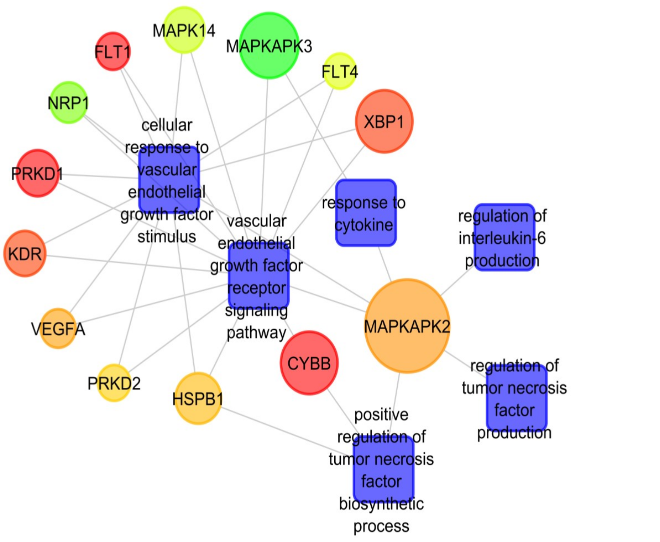


**c)**

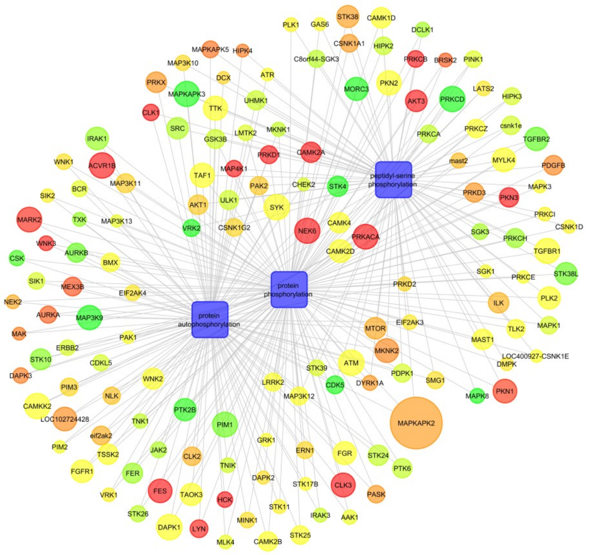

**d)**

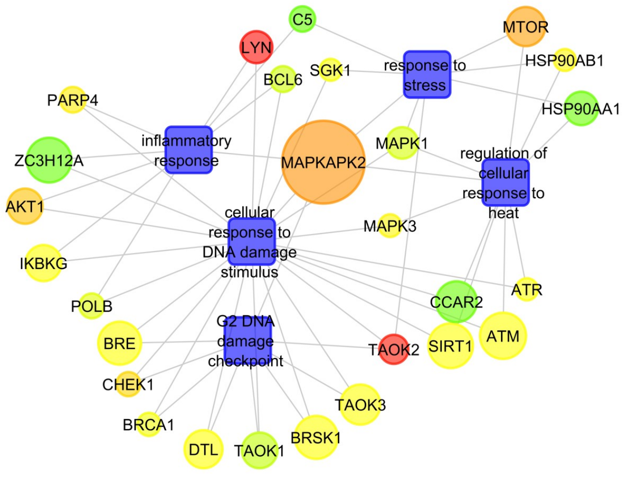

**e)**

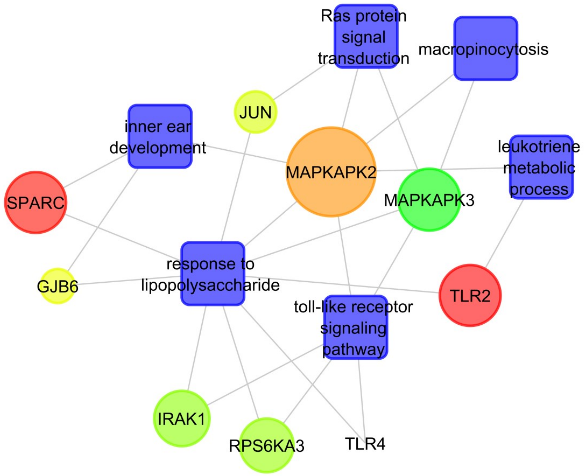


**Figure S8:** Clustering of the major biological pathways regulated by MK2 (F vs E group). The diagrammatic representation of the transcriptome profiling data keeping MK2 at the nexus of the analysis portrays that it is playing a major role in the regulation of six major biological processes, out of which five are as shown. **a)** MAPK-Pathway **b)** Regulation of Cytokines **c)** Protein phosphorylation **d)** Response to stress, and **e)** Signal transduction

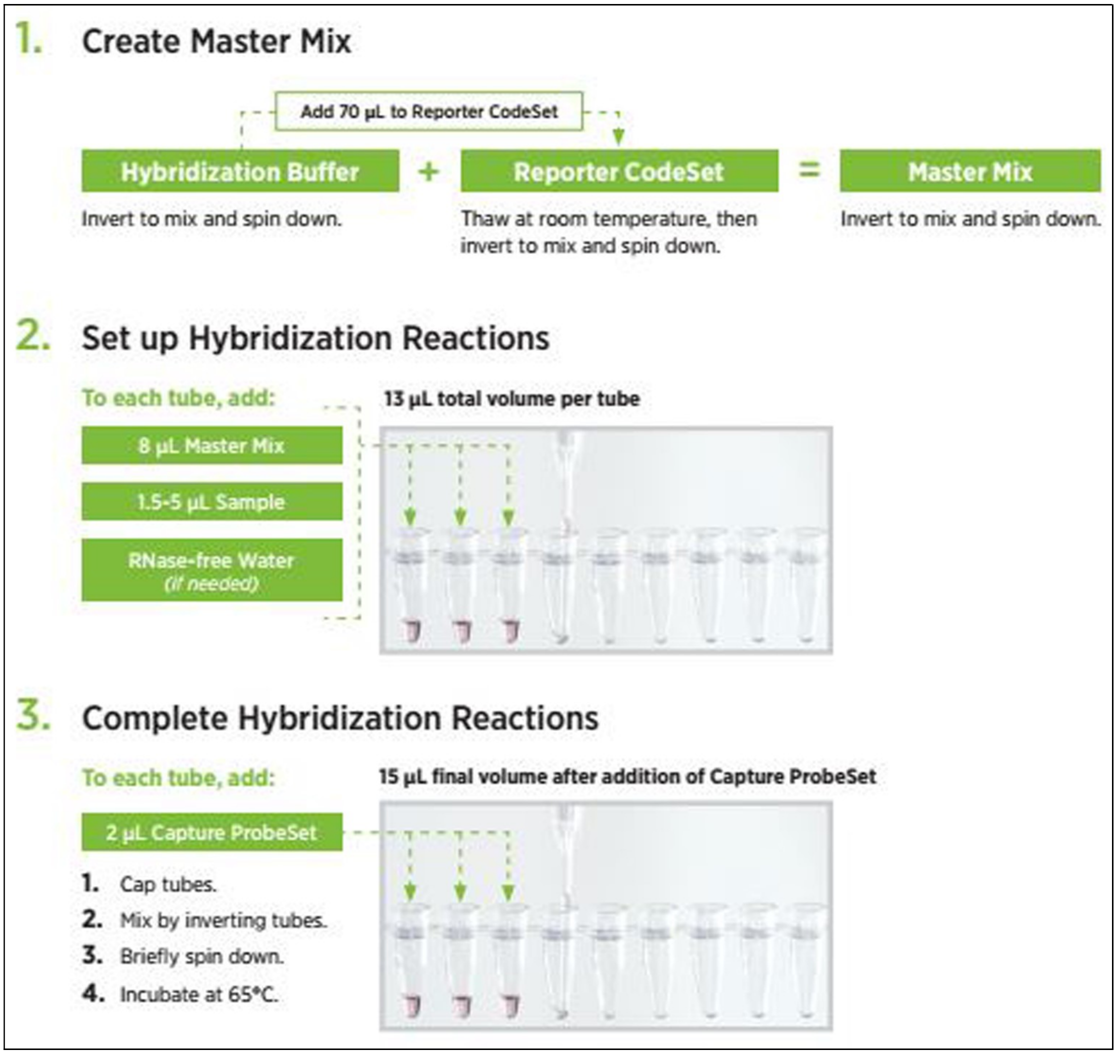


**Figure S9:** Schematic workflow of hybridization process in the nCounter gene expression assay showing a stepwise detailing of the approach employed for the assay.
