## Supplemental Tables for "RNA-sequencing manifests the intrinsic role of MAPKAPK2 in facilitating molecular crosstalk during HNSCC pathogenesis"

### Additional File 2

**Authors' information**

[1]Pharmacology and Toxicology Laboratory, CSIR-Institute of Himalayan Bioresource Technology (CSIR-IHBT), Palampur-176061, India

[2]Biotechnology Division, CSIR-Institute of Himalayan Bioresource Technology (CSIR-IHBT), Palampur-176061, India

[3]Academy of Scientific and Innovative Research (AcSIR), Ghaziabad-201002, India.

**Tables:**

**Table ST1: List of antibodies used for IHC analysis**

| S. No. | Antibody | Catalog Number | Dilution Used | Raised in | Make |
|---|---|---|---|---|---|
| 1 | MELK | MA-517120 | 1:200 | Mouse | THERMO |
| 2 | ZNF662 | PA5-58662 | 1:200 | Rabbit | THERMO |
| 3 | CREB3L1 | PA5-13537 | 1:50 | Rabbit | THERMO |
| 4 | IGFBP2 | MA5-15400 | 1:200 | Mouse | THERMO |
| 5 | MUC4 | 35-4900 | 1:50 | Mouse | THERMO |
| 6 | PRKARB2 | PA5-28266 | 1:100 | Rabbit | THERMO |
| 7 | BMP-7 | MA5-23878 | 1:20 | Mouse | THERMO |

**Table ST2:** Tabular representation of the list of the 77 common elements/biological processes in the transcriptome profiling of the *in vitro* HNSCC cell line model (A-D comparisons).

| S. No. | Biological Processes |
|---|---|
| 1. | Alternative splicing |
| 2. | Sequence variant |
| 3. | Polymorphism |
| 4. | Phosphoprotein |
| 5. | GO:0005515~protein binding |
| 6. | Splice variant |
| 7. | GO:0005737~cytoplasm |
| 8. | GO:0005634~nucleus |
| 9. | Cytoplasm |
| 10. | Nucleus |
| 11. | GO:0005829~cytosol |
| 12. | Acetylation |
| 13. | Metal-binding |
| 14. | Coiled coil |
| 15. | GO:0005654~nucleoplasm |
| 16. | GO:0016020~membrane |
| 17. | Mutagenesis site |
| 18. | Ubl conjugation |
| 19. | Zinc |
| 20. | Transcription |
| 21. | Transcription regulation |
| 22. | Nucleotide-binding |
| 23. | Isopeptide bond |
| 24. | GO:0005524~ATP binding |
| 25. | ATP-binding |
| 26. | Cytoskeleton |
| 27. | GO:0005794~Golgi apparatus |
| 28. | GO:0044822~poly(A) RNA binding |
| 29. | GO:0045944~positive regulation of transcription from RNA polymerase II promoter |
| 30. | Compositionally biased region:Pro-rich |
| 31. | Methylation |
| 32. | GO:0000122~negative regulation of transcription from RNA polymerase II promoter |
| 33. | GO:0043231~intracellular membrane-bounded organelle |
| 34. | Cell junction |
| 35. | Apoptosis |
| 36. | GO:0048471~perinuclear region of cytoplasm |
| 37. | Activator |
| 38. | GO:0006915~apoptotic process |
| 39. | Repressor |
| 40. | Compositionally biased region:Ser-rich |

| | |
|---|---|
| 41. | GO:0005913~cell-cell adherens junction |
| 42. | RNA-binding |
| 43. | GO:0006468~protein phosphorylation |
| 44. | GO:0005925~focal adhesion |
| 45. | GO:0098641~cadherin binding involved in cell-cell adhesion |
| 46. | GO:0098609~cell-cell adhesion |
| 47. | GO:0019901~protein kinase binding |
| 48. | Host-virus interaction |
| 49. | GO:0003682~chromatin binding |
| 50. | Chromatin regulator |
| 51. | Chromosomal rearrangement |
| 52. | GO:0005856~cytoskeleton |
| 53. | Short sequence motif:Nuclear localization signal |
| 54. | IPR016024:Armadillo-type fold |
| 55. | Ligase |
| 56. | Serine/threonine-protein kinase |
| 57. | GO:0015629~actin cytoskeleton |
| 58. | Actin-binding |
| 59. | GO:0016874~ligase activity |
| 60. | GO:0008134~transcription factor binding |
| 61. | GO:0005096~GTPase activator activity |
| 62. | GTPase activation |
| 63. | GO:0005516~calmodulin binding |
| 64. | GO:0005815~microtubule organizing center |
| 65. | GO:0005938~cell cortex |
| 66. | Calmodulin-binding |
| 67. | IPR016201:Plexin-like fold |
| 68. | SM00423:PSI |
| 69. | IPR019787:Zinc finger, PHD-finger |
| 70. | IPR001589:Actinin-type, actin-binding, conserved site |
| 71. | Domain:CH 1 |
| 72. | Domain:CH 2 |
| 73. | Repeat:Spectrin 4 |
| 74. | Repeat:Spectrin 3 |
| 75. | Repeat:Spectrin 2 |
| 76. | Repeat:Spectrin 1 |
| 77. | Domain:CNH |

**Table ST3:** Tabular representation of list of the five common genes in the 77 common elements in the transcriptome profiling of the *in vitro* HNSCC cell line model (A-D comparisons). The average log2 fold change values are colored according to the change in gene expression (red indicates upregulation while green indicates downregulation).

| 5 common elements | B_Vs_A | D_Vs_C | D_Vs_B | C_Vs_A | Gene Name | OFFICIAL_GENE_SYMBOL |
|---|---|---|---|---|---|---|
| NM_001142614 | -4.74 | 3.66 | 5.16 | -3.28 | EH domain binding protein 1(EHBP1) | EHBP1 |
| NM_018121 | -7.09 | 6.44 | 6.47 | -7.10 | SMC5-SMC6 complex localization factor 2(SLF2) | SLF2 |
| NM_033657 | 7.68 | -3.85 | -3.95 | 7.54 | death associated protein 3(DAP3) | DAP3 |
| XM_006713202 | 7.72 | -7.91 | -7.75 | 7.84 | inositol hexakisphosphate kinase 2(IP6K2) | IP6K2 |
| XM_011529767 | -5.94 | 6.28 | 5.58 | -6.68 | runt related transcription factor 1(RUNX1) | RUNX1 |

**Table ST4:** Tabular representation of list of the 16 common elements/biological processes in the transcriptome profiling of the *in vitro* HNSCC cell line model (A-D comparisons) filtered out on the basis of relevance in cancer progression.

| S. No. | Biological Processes |
|---|---|
| 1. | Transcription |
| 2. | Transcription regulation |
| 3. | Cytoskeleton |
| 4. | GO:0044822~poly(A) RNA binding |
| 5. | GO:0045944~positive regulation of transcription from RNA polymerase II promoter |
| 6. | GO:0000122~negative regulation of transcription from RNA polymerase II promoter |
| 7. | Apoptosis |
| 8. | GO:0006915~apoptotic process |
| 9. | RNA-binding |
| 10. | GO:0006468~protein phosphorylation |
| 11. | GO:0098609~cell-cell adhesion |
| 12. | GO:0019901~protein kinase binding |
| 13. | GO:0005856~cytoskeleton |
| 14. | Serine/threonine-protein kinase |
| 15. | GO:0015629~actin cytoskeleton |
| 16. | GO:0008134~transcription factor binding |

**Table ST5:** Tabular representation of the list of the two common genes in the 16 common elements in the transcriptome profiling of the *in vitro* HNSCC cell line model (A-D comparisons). The average log2 fold change values are colored according to the change in gene expression (red indicates upregulation while green indicates downregulation).

| 2 common elements | B_Vs_A | D_Vs_C | D_Vs_B | C_Vs_A | Gene Name | OFFICIAL_GENE_SYMBOL |
|---|---|---|---|---|---|---|
| NM_033657 | 7.68 | -3.85 | -3.95 | 7.54 | death associated protein 3(DAP3) | DAP3 |
| XM_011529767 | -5.94 | 6.28 | 5.58 | -6.68 | runt related transcription factor 1(RUNX1) | RUNX1 |

**Table ST6:** Tabular representation of list of the 14 common elements/biological processes in the transcriptome profiling of the *in vivo* heterotopic HNSCC xenograft experimental dataset (F vs E comparison) filtered out on the basis of relevance in cancer progression.

| S. No. | Biological Processes |
|---|---|
| 1. | Transcription |
| 2. | Transcription regulation |
| 3. | Cytoskeleton |
| 4. | GO:0045944~positive regulation of transcription from RNA polymerase II promoter |
| 5. | GO:0044822~poly(A) RNA binding |
| 6. | GO:0000122~negative regulation of transcription from RNA polymerase II promoter |
| 7. | RNA-binding |
| 8. | GO:0006468~protein phosphorylation |
| 9. | Apoptosis |
| 10. | Serine/threonine-protein kinase |
| 11. | GO:0019901~protein kinase binding |
| 12. | GO:0008134~transcription factor binding |
| 13. | GO:0098609~cell-cell adhesion |
| 14. | GO:0015629~actin cytoskeleton |

**Table ST7:** Tabular representation of the list of the top two up- and down regulated genes in selected 16 cancer specific pathways in the transcriptome profiling of the *in vitro* HNSCC cell line model (A-D comparisons) following 3'-untranslated region-based filtering.

| S. No. | Gene Symbol | Entrez Gene ID |
|---|---|---|
| 1. | CALD1 | 800 |
| 2. | CLK2 | 1196 |
| 3. | CTBP2 | 1488 |
| 4. | ERF | 2077 |
| 5. | FER | 2241 |
| 6. | NR3C1 | 2908 |
| 7. | IRAK1 | 3654 |
| 8. | PPP1R12A | 4659 |
| 9. | BRD2 | 6046 |
| 10. | SNAPC4 | 6621 |
| 11. | SP3 | 6670 |
| 12. | TAF1 | 6872 |
| 13. | UTRN | 7402 |
| 14. | ZNF189 | 7743 |
| 15. | KDM5C | 8242 |
| 16. | LATS1 | 9113 |
| 17. | MAP4K4 | 9448 |
| 18. | H2AFY | 9555 |
| 19. | NCOR1 | 9611 |
| 20. | MELK | 9833 |
| 21. | PAK4 | 10298 |
| 22. | CAMKK2 | 10645 |
| 23. | ADNP2 | 22850 |
| 24. | FOXJ3 | 22887 |
| 25. | NCOA6 | 23054 |
| 26. | CAMTA2 | 23125 |
| 27. | PASK | 23178 |
| 28. | DICER1 | 23405 |
| 29. | BRD4 | 23476 |
| 30. | BAZ2B | 29994 |
| 31. | MINK1 | 50488 |
| 32. | SAMD4B | 55095 |
| 33. | PBRM1 | 55193 |
| 34. | NEK9 | 91754 |

**Table ST8:** Tabular representation of list of the topmost up- and down regulated genes in cancer specific pathways in the transcriptome profiling of the *in vivo* heterotopic HNSCC xenograft experimental dataset (F vs E comparison) following 3'-untranslated region-based filtering.

| S. No. | Gene Symbol | Entrez Gene ID |
|---|---|---|
| 1. | APBB2 | 323 |
| 2. | BMP7 | 655 |
| 3. | DST | 667 |
| 4. | DDR1 | 780 |
| 5. | CDC25B | 994 |
| 6. | CDSN | 1041 |
| 7. | DPYSL3 | 1809 |
| 8. | EIF4E | 1977 |
| 9. | EZH1 | 2145 |
| 10. | FOXO3 | 2309 |
| 11. | GUK1 | 2987 |
| 12. | HNRNPD | 3184 |
| 13. | IGFBP2 | 3485 |
| 14. | ITPR1 | 3708 |
| 15. | JAK1 | 3716 |
| 16. | LIMK1 | 3984 |
| 17. | LIMS1 | 3987 |
| 18. | SMAD3 | 4088 |
| 19. | MUC4 | 4585 |
| 20. | PFKM | 5213 |
| 21. | PFKP | 5214 |
| 22. | PI4KB | 5298 |
| 23. | PPP2R1B | 5519 |
| 24. | PRKAR2B | 5577 |
| 25. | SS18 | 6760 |
| 26. | UBE3A | 7337 |
| 27. | ZNF131 | 7690 |
| 28. | SEMA7A | 8482 |
| 29. | ZBED1 | 9189 |
| 30. | TGFBRAP1 | 9392 |
| 31. | H2AFY | 9555 |
| 32. | STK25 | 10494 |
| 33. | TRAK1 | 22906 |
| 34. | ZC3H13 | 23091 |
| 35. | ATP13A2 | 23400 |
| 36. | PTPN22 | 26191 |
| 37. | ZNF544 | 27300 |
| 38. | PPP1R12C | 54776 |
| 39. | PCBP4 | 57060 |

| | | |
|---|---|---|
| 40. | NDRG2 | 57447 |
| 41. | MKL2 | 57496 |
| 42. | KMT2C | 58508 |
| 43. | CREB3L1 | 90993 |
| 44. | CPEB2 | 132864 |
| 45. | FREM1 | 158326 |
| 46. | ZNF48 | 197407 |
| 47. | SLC35B2 | 347734 |
| 48. | ZNF662 | 389114 |

**Table ST9:** Tabular representation of the CodeSet details showing all the information of the molecular probes that were used for the nCounter gene expression assay.

**CODESET DETAILS**

| | Customer Identifier | Accession | Position | Target Sequence | Tm CP | Tm RP | Flags | HUGO Gene | N SID |
|---|---|---|---|---|---|---|---|---|---|
| 1 | ABCF1 | NM_001090.2 | 851-950 | GATGTCCTCCCGCCA | 79 | 82 | HK | ABCF1 | NM_001090.2:850 |
| 2 | ADNP2 | NM_014913.3 | 1076-1175 | GAAGATCCCACCAC | 83 | 80 | | ADNP2 | NM_014913.3:1075 |
| 3 | APBB2 | NM_004307.1 | 2418-2517 | GGGCATGTTACCTGT | 81 | 84 | | APBB2 | NM_004307.1:2417 |
| 4 | ATP13A2 | NM_001141974.1 | 926-1025 | TGGCTGGCTGACCAC | 83 | 84 | | ATP13A2 | NM_001141974.1:925 |
| 5 | BAZ2B | NM_013450.2 | 3901-4000 | TATGGAAGCCCACTG | 85 | 83 | | BAZ2B | NM_013450.2:3900 |
| 6 | BMP7 | NM_001719.1 | 526-625 | GCTTCGTCAACCTCG | 79 | 79 | | BMP7 | NM_001719.1:525 |
| 7 | BRD2 | NM_005104.2 | 1891-1990 | CCGGAGGTGTCCAA | 83 | 79 | | BRD2 | NM_005104.2:1890 |
| 8 | BRD4 | NM_014299.2 | 746-845 | AGGCAAAAGGAAGA | 83 | 82 | | BRD4 | NM_014299.2:745 |
| 9 | CALD1 | NM_004342.6 | 3863-3962 | GAAGTCAAACCCTCA | 83 | 78 | | CALD1 | NM_004342.6:3862 |
| 10 | CAMKK2 | NM_006549.3 | 1711-1810 | GGATCTGATCAAAGG | 81 | 82 | | CAMKK2 | NM_006549.3:1710 |
| 11 | CAMTA2 | NM_015099.2 | 1574-1673 | TTGGAGGACCTGTTG | 83 | 83 | | CAMTA2 | NM_015099.2:1573 |
| 12 | CDC25B | NM_021873.2 | 3046-3145 | CACCATACGAGCAC | 80 | 78 | | CDC25B | NM_021873.2:3045 |
| 13 | CDSN | NM_001264.4 | 965-1064 | TGACAGTTATCTGGT | 83 | 83 | | CDSN | NM_001264.4:964 |
| 14 | CLK2 | NM_003993.2 | 903-1002 | CGACTTGAGATCAAC | 83 | 83 | | CLK2 | NM_003993.2:902 |
| 15 | CPEB2 | NM_001177381.1 | 2562-2661 | GGATCCCCGAAAAA | 79 | 80 | | CPEB2 | NM_001177381.1:2561 |
| 16 | CREB3L1 | NM_052854.1 | 196-295 | CAACAATGCGCACTT | 83 | 82 | | CREB3L1 | NM_052854.1:195 |
| 17 | CTBP2 | NM_001329.2 | 1885-1984 | GTTGCAGTGTTGAAA | 82 | 80 | | CTBP2 | NM_001329.2:1884 |
| 18 | DAP3 | NM_004632.3 | 796-895 | CACAGATGCAGTTGG | 82 | 82 | | DAP3 | NM_004632.3:795 |
| 19 | DDR1 | NM_001954.4 | 1343-1442 | CCCTTGGCGGCCGT | 83 | 82 | | DDR1 | NM_001954.4:1342 |
| 20 | DICER1 | NM_177438.2 | 5061-5160 | AGAATTTCAACAGCC | 83 | 79 | | DICER1 | NM_177438.2:5060 |
| 21 | DPYSL3 | NM_001197294.1 | 927-1026 | GAAGAGTTGCTGTGA | 84 | 84 | | DPYSL3 | NM_001197294.1:926 |
| 22 | DST | NM_001723.4 | 1871-1970 | CGAATAACTCTCATC | 80 | 84 | | DST | NM_001723.4:1870 |
| 23 | EHBP1 | NM_015252.3 | 1751-1850 | AGATCTCTCTACTTCT | 80 | 79 | | EHBP1 | NM_015252.3:1750 |
| 24 | EIF4E | NM_001130678.1 | 594-693 | TGGAGAATCTTTTGA | 77 | 81 | | EIF4E | NM_001130678.1:593 |
| 25 | ERF | NM_006494.2 | 2008-2107 | TCCCTGTCTCTGTGG | 86 | 84 | | ERF | NM_006494.2:2007 |
| 26 | EZH1 | NM_001991.3 | 2721-2820 | CAACTTAGGCAGTTC | 81 | 81 | | EZH1 | NM_001991.3:2720 |
| 27 | FER | NM_005246.2 | 2002-2101 | AAGAAATCAGGTGTA | 80 | 82 | | FER | NM_005246.2:2001 |
| 28 | FOXJ3 | NM_014947.3 | 2221-2320 | TACTGCTTTTATCTGA | 79 | 79 | | FOXJ3 | NM_014947.3:2220 |
| 29 | FOXO3 | NM_001455.2 | 1861-1960 | CCGGAACGTGATGC | 82 | 80 | X | FOXO3 | NM_001455.2:1860 |
| 30 | FREM1 | NM_001177704.1 | 615-714 | GGGCTCCTGAAAAA | 84 | 82 | | FREM1 | NM_001177704.1:614 |
| 31 | GAPDH | NM_001256799.1 | 387-486 | GAACGGGAAGCTTG | 85 | 86 | HK | GAPDH | NM_001256799.1:386 |
| 32 | GUK1 | NM_000858.5 | 431-530 | CGAGGCCCGGCGAG | 82 | 83 | | GUK1 | NM_000858.5:430 |
| 33 | H2AFY | NM_001040158.1 | 546-645 | GTTGCTAGCGAAGA | 79 | 83 | | H2AFY | NM_001040158.1:545 |
| 34 | HNRNPD | NM_002138.3 | 695-794 | GCACTCTGAAGTTAG | 81 | 78 | | HNRNPD | NM_002138.3:694 |
| 35 | IGFBP2 | NM_000597.2 | 676-775 | TCGGGTATGAAGGA | 85 | 85 | | IGFBP2 | NM_000597.2:675 |
| 36 | IP6K2 | NM_001005910.2 | 278-377 | ATGTGGAGCCCCGC | 85 | 86 | | IP6K2 | NM_001005910.2:277 |
| 37 | IRAK1 | NM_001569.3 | 1996-2095 | CACAGCCGTGGAAG | 82 | 82 | | IRAK1 | NM_001569.3:1995 |
| 38 | ITPR1 | NM_001099952.2 | 4819-4918 | TCTTCAGCTCTCCCTT | 83 | 82 | | ITPR1 | NM_001099952.2:4818 |
| 39 | JAK1 | NM_002227.1 | 286-385 | GAGAACACCAAGCT | 81 | 79 | | JAK1 | NM_002227.1:285 |
| 40 | KDM5C | NM_004187.2 | 1171-1270 | CCAAGAGACTGCAG | 82 | 82 | | KDM5C | NM_004187.2:1170 |
| 41 | KMT2C | NM_170606.2 | 11974-12073 | GGTTTTGCAACAACT | 83 | 81 | | KMT2C | NM_170606.2:11973 |
| 42 | LATS1 | NM_004690.2 | 1091-1190 | TAAAGAATCCTTAGT | 81 | 82 | | LATS1 | NM_004690.2:1090 |
| 43 | LIMK1 | NM_002314.3 | 3134-3233 | GGGATTTTATTTTTGT | 80 | 78 | | LIMK1 | NM_002314.3:3133 |
| 44 | LIMS1 | NM_004987.3 | 1409-1508 | AAAGGAGTCACCTTT | 76 | 77 | | LIMS1 | NM_004987.3:1408 |
| 45 | MAP4K4 | NM_004834.3 | 3316-3415 | GATGCCTACATCAGT | 81 | 81 | | MAP4K4 | NM_004834.3:3315 |
| 46 | MELK | NM_014791.2 | 801-900 | TATGAGAGGAAAATA | 82 | 81 | | MELK | NM_014791.2:800 |
| 47 | MINK1 | NM_001024937.1 | 3346-3445 | CCTGCTCATCACCAT | 80 | 82 | | MINK1 | NM_001024937.1:3345 |
| 48 | MKL2 | NM_014048.3 | 1786-1885 | TGAGAATGACAAATA | 81 | 80 | | MRTFB | NM_014048.3:1785 |
| 49 | MUC4 | NM_018406.4 | 14311-14410 | CAGCCTGGGCCCCG | 83 | 83 | | MUC4 | NM_018406.4:14310 |
| 50 | NCOA6 | NM_014071.3 | 1948-2047 | GAATTCAGGAGCCC | 85 | 85 | | NCOA6 | NM_014071.3:1947 |
| 51 | NCOR1 | NM_006311.3 | 1391-1490 | TAGGAGTGAGCATG | 82 | 83 | | NCOR1 | NM_006311.3:1390 |
| 52 | NDRG2 | NM_016250.2 | 1516-1615 | TATGCATCCTCTGTC | 82 | 81 | | NDRG2 | NM_016250.2:1515 |
| 53 | NEK9 | NM_033116.3 | 2086-2185 | GGCTCTGATATCTGT | 81 | 81 | | NEK9 | NM_033116.3:2085 |
| 54 | NR3C1 | NM_001018077.1 | 2823-2922 | GGAGATCATATAGAC | 80 | 81 | | NR3C1 | NM_001018077.1:2822 |
| 55 | PAK4 | NM_005884.3 | 2131-2230 | CCTCCTTGCTGGGGG | 82 | 82 | | PAK4 | NM_005884.3:2130 |
| 56 | PASK | NM_015148.3 | 3233-3332 | AAGGTCTTGGAGGA | 82 | 81 | | PASK | NM_015148.3:3232 |
| 57 | PBRM1 | NM_018313.4 | 207-306 | AAGCAGGAAAAGGA | 81 | 83 | | PBRM1 | NM_018313.4:206 |
| 58 | PCBP4 | NM_020418.2 | 811-910 | CATCACCGGGTCTAC | 82 | 79 | | PCBP4 | NM_020418.2:810 |
| 59 | PFKM | NM_000289.5 | 2196-2295 | TTTGATAGGAATTTTG | 83 | 81 | | PFKM | NM_000289.5:2195 |
| 60 | PFKP | NM_002627.3 | 796-895 | GTCAAACTCTCGGAG | 81 | 84 | | PFKP | NM_002627.3:795 |
| 61 | PI4KB | NM_001198775.1 | 2711-2810 | TGCTTTTTCCCTGCCT | 83 | 81 | | PI4KB | NM_001198775.1:2710 |
| 62 | POLR2A | NM_000937.2 | 3776-3875 | TTCCAAGAAGCCAAA | 78 | 84 | HK | POLR2A | NM_000937.2:3775 |
| 63 | PPP1R12A | NM_001143886.1 | 351-450 | AAGTTAATCGGCAAG | 83 | 83 | | PPP1R12A | NM_001143886.1:350 |
| 64 | PPP1R12C | NM_001271618.1 | 2333-2432 | CTGACAACCAGCGC | 83 | 84 | | PPP1R12C | NM_001271618.1:2332 |
| 65 | PPP2R1B | NM_002716.4 | 1446-1545 | GCTGAATTCTTTATGT | 80 | 84 | | PPP2R1B | NM_002716.4:1445 |
| 66 | PRKAR2B | NM_002736.2 | 1351-1450 | GAAAAGGAACATCG | 81 | 83 | | PRKAR2B | NM_002736.2:1350 |
| 67 | PTPN22 | NM_012411.5 | 343-442 | CCGGGTAGAACTATC | 82 | 81 | | PTPN22 | NM_012411.5:342 |
| 68 | RPL19 | NM_000981.3 | 316-415 | CCAATGCCCGAATG | 84 | 81 | HK;X | RPL19 | NM_000981.3:315 |
| 69 | RUNX1 | NM_001754.4 | 636-735 | CAGCCATGAAGAAC | 80 | 83 | | RUNX1 | NM_001754.4:635 |
| 70 | SAMD4B | NM_018028.2 | 1337-1436 | ACATGAGGCTACTGC | 83 | 83 | | SAMD4B | NM_018028.2:1336 |
| 71 | SEMA7A | NM_001146029.1 | 661-760 | CCCACAGTTCATCAA | 83 | 83 | | SEMA7A | NM_001146029.1:660 |
| 72 | SLC35B2 | NM_178148.3 | 1185-1284 | CTGCTACTCTCCATC | 81 | 83 | | SLC35B2 | NM_178148.3:1184 |
| 73 | SLF2 | NM_001243770.1 | 254-353 | CCCCGCCTTCTCCAG | 85 | 85 | | SLF2 | NM_001243770.1:253 |
| 74 | SMAD3 | NM_005902.3 | 4221-4320 | TTAAAGGACAGTTGA | 79 | 80 | | SMAD3 | NM_005902.3:4220 |
| 75 | SNAPC4 | NM_003086.2 | 1160-1259 | CCATGCAGCTGATCT | 85 | 86 | | SNAPC4 | NM_003086.2:1159 |
| 76 | SP3 | NM_003111.3 | 656-755 | AGTGTCCAGTGTTCA | 81 | 79 | | SP3 | NM_003111.3:655 |
| 77 | SS18 | NM_001007559.1 | 367-466 | CCACCTCCACGCTCT | 85 | 86 | | SS18 | NM_001007559.1:366 |
| 78 | STK25 | NM_006374.3 | 1431-1530 | GTGCACCTGGTGGA | 82 | 82 | | STK25 | NM_006374.3:1430 |
| 79 | TAF1 | NM_004606.4 | 1067-1166 | GAAATCACGATGATG | 81 | 83 | | TAF1 | NM_004606.4:1066 |
| 80 | TGFBRAP1 | NM_001142621.1 | 1586-1685 | ATGCTGCTGCAGTTC | 83 | 79 | | TGFBRAP1 | NM_001142621.1:1585 |
| 81 | TRAK1 | NM_014965.3 | 1355-1454 | CGGTCCAGCTTCTAC | 85 | 85 | | TRAK1 | NM_014965.3:1354 |
| 82 | UBE3A | NM_000462.2 | 2736-2835 | CCAGTACAAAAAGG | 82 | 83 | | UBE3A | NM_000462.2:2735 |
| 83 | UTRN | NM_007124.2 | 8387-8486 | TCCCTCTCTAAAGAT | 83 | 79 | | UTRN | NM_007124.2:8386 |
| 84 | ZBED1 | NM_001171136.1 | 4086-4185 | TTTAAGTCTGTGGAA | 82 | 81 | | ZBED1 | NM_001171136.1:4085 |
| 85 | ZC3H13 | NM_015070.3 | 3896-3995 | AAGAAGAAGTTCTCC | 83 | 82 | | ZC3H13 | NM_015070.3:3895 |
| 86 | ZNF131 | NM_003432.1 | 623-722 | CAAGCAGTCCGTAA | 81 | 83 | | ZNF131 | NM_003432.1:622 |
| 87 | ZNF189 | NM_197977.2 | 2131-2230 | TCAACAGCGCAGTCT | 83 | 82 | | ZNF189 | NM_197977.2:2130 |
| 88 | ZNF48 | NM_001214907.1 | 2585-2684 | AGAGACAGAAGGTG | 84 | 84 | | ZNF48 | NM_001214907.1:2584 |
| 89 | ZNF544 | NM_014480.3 | 283-382 | TCCTGGCACAGCGG | 92 | 80 | | ZNF544 | NM_014480.3:282 |
| 90 | ZNF662 | NM_207404.3 | 715-814 | GGAATTATGGTTTGG | 84 | 82 | | ZNF662 | NM_207404.3:714 |

**Table ST10:** List of housekeeping genes used in nCounter expression assay.

| S. No. | Housekeeping Gene | Entrez Gene ID |
|---|---|---|
| 1. | GAPDH | 2597 |
| 2. | POLR2A | 5430 |
| 3. | ABCF1 | 23 |
| 4. | RPL19 | 6143 |
